# Ontology-Aware Deep Learning Enables Ultrafast, Accurate and Interpretable Source Tracking among Sub-Million Microbial Community Samples from Hundreds of Niches

**DOI:** 10.1101/2020.11.01.364208

**Authors:** Yuguo Zha, Hui Chong, Hao Qiu, Kai Kang, Yuzheng Dun, Zhixue Chen, Xuefeng Cui, Kang Ning

## Abstract

The taxonomical structure of microbial community sample is highly habitat-specific, making it possible for source tracking niches where samples are originated. Current methods face challenges when the number of samples and niches are magnitudes more than current in use, under which circumstances they are unable to accurately source track samples in a timely manner, rendering them difficult in knowledge discovery from sub-million heterogeneous samples. Here, we introduce a deep learning method based on Ontology-aware Neural Network approach, ONN4MST (https://github.com/HUST-NingKang-Lab/ONN4MST), which takes into consideration the ontology structure of niches and the relationship of samples from these ontologically-organized niches. ONN4MST’s superiority in accuracy, speed and robustness have been proven, for example with an accuracy of 0.99 and AUC of 0.97 in a microbial source tracking experiment that 125,823 samples and 114 niches were involved. Moreover, ONN4MST has been utilized on several source tracking applications, showing that it could provide highly-interpretable results from samples with previously less-studied niches, detect microbial contaminants, and identify similar samples from ontologically-remote niches, with high fidelity.

## Introduction

With the rapid accumulation of microbial community samples from various niches (biomes) around the world, as well as the huge volume of sequencing data deposited into public databases, such as those from the “Human Microbiome Project”^1,2^ and the “Earth Microbiome Project”^3,4^, knowledge about microbial communities and their influence on environment and human health has grown rapidly^5,6^. Such massive amount of microbial community samples provide the opportunity to study the inconspicuous evolution and ecological patterns among microbial communities, especially habitat-specific patterns.

One key challenge faced with such a paramount number of heterogeneous samples is to track potential origin of a microbial community sample, as well as distinguishing samples from different health conditions or diverse environments, calling for fast and accurate source tracking^7–9^. Taxonomical composition of a microbial community sample is usually represented by hierarchically-structured taxa and their relative abundances (also referred to as the community structure), and these taxa are functioning in concert to maintain the stability of the microbial community and its adaptation to the specific environment (also referred to as the niche or biome)^10,11^. Biomes are organized in an ontology structure with six different layers (simply referred to as the biome ontology). Layer one is the highest layer containing only one biome “Root” and layer six is the lowest (bottom) layer containing biomes such as “Fecal”. Biomes of lower layers such as “Human gut” belong to those of higher layers such as “Human digestive system”, whereby EBI MGnify currently contain the most up-to-date biome structure^11^ with more than one hundred biomes as of January 2020. In general, microbial community samples from the same biome tend to have similar community structures, while such similarities are highly dependent on the biome layers. Source tracking the microbial community samples, especially among a massive amount of samples, remains a challenging problem today.

Several methods for microbial community source tracking have already been proposed^9,12–15^. They can generally be divided into two categories, namely distance-based methods such as Jensen-Shannon Divergence (JSD)^16^, Striped UniFrac^13^ and Meta-Prism^17^, and unsupervised machine learning methods such as SourceTracker based on Bayesian algorithm^15^ and FEAST based on Expected-Maximization algorithm^9^. However, the limitations of these methods are apparent: Firstly, due to the nature of distance-based method and unsupervised method, they are relatively slow, especially when the number of samples exceed tens of thousands^9^, hindering them from identifying potential source environments in a timely manner. Secondly, there is still a lack of method for accurate source tracking from more than a hundred biomes, largely due to the resolution limitation of both methods^9,15^. Thirdly, current methods are not suitable for knowledge discovery of samples from previously less studied or unknown biomes.

To address these limitations, we developed ONN4MST, an Ontology-aware Neural Network (ONN) computational model for microbial source tracking. It is a supervised learning method, and has utilized the biome ontology information. It has provided an ultrafast (less than 0.1 seconds) and accurate (AUC higher than 0.97 in most cases) solution for searching a sample against dataset containing more than a hundred potential biomes and sub-million samples, and also out-performed state-of-the-art methods in scalability and stability. The ability of ONN4MST on knowledge discovery is also demonstrated by utilization in various source tracking applications: it enables source tracking of samples whose niches are previously less studied or unknown, detection of microbial contaminants, as well as identification of similar samples from ontologically-remote biomes, showing the unique importance of ONN4MST in knowledge discovery from huge amount of microbial community samples of heterogeneous biomes.

## Results

### Ontology-aware Neural Network

ONN4MST uses an Ontology-aware Neural Network (ONN) model for source tracking. When training the model, all training samples’ community structures are decoded, each converted to a matrix containing the taxa at different taxonomical levels and their relative abundances (simply referred to as the Matrix). The ONN model uses the Matrix as input and reshapes it into tensors which point to biomes at every different layer of the biome ontology. To fit the structure of biome ontology, the ONN model uses multiple ontology units, each belonging to one of the six specific layers of biome ontology (**Fig. 1a**). The conceptual modules, the training procedure and the evaluation procedure of the ONN model are illustrated in **Supplementary Fig. 1** and described in **Methods**.

**Fig. 1:**
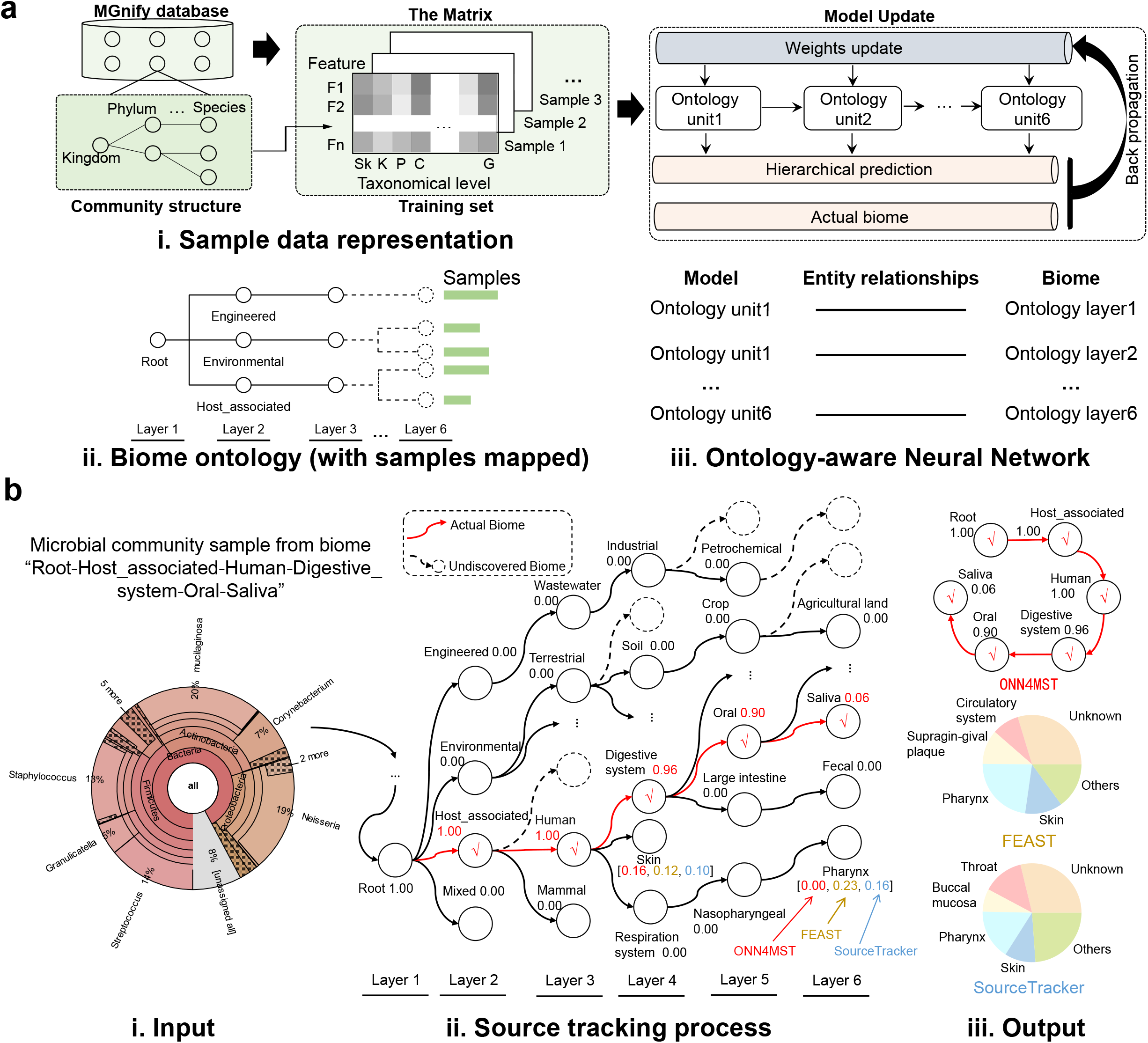
Building and using the Ontology-aware Neural Network model for microbial source tracking. **a**. The sample data representation and training process of ONN model. **i**. Sample data are transformed into the Matrix. With the Matrix, each column represents a taxonomical level and each row represents a feature; **ii.** In parallel, samples are mapped to biome ontology according to their niches; **iii**. The model is built and updated according to both samples’ abundance matrices and biome ontology information. More details about building, testing and using the ONN model for source tracking are illustrated in **Supplementary Fig. 1** and **Supplementary Fig. 2. b**. An illustrated example of microbial source tracking procedure using ONN4MST. **i**. The input is the community structure of a real microbial community sample (this sample is from the biome “Root-Host_associated-Human-Digestive_system-Oral-Saliva”) that has been preprocessed and the Matrix has been provided into the model; **ii.** Source tracking process at different layers. The red arrows indicate the search process from layer 1 to layer 6, accompanied with source contribution annotated in red. To compare with the procedure of ONN4MST, the yellow and blue arrows indicated the source tracking results (among the overall top 5 sources) of FEAST and Source Tracker, together with their source contributions, respectively. The actual biome is annotated by a red check mark; **iii.** The predicted biomes (with source contributions) by ONN4MST, FEAST and SourceTracker.

The source tracking procedure of ONN4MST is illustrated in **Fig. 1b**. Since ONN4MST is the first method available that could source track the samples at different layers of biome ontology, the search scheme of ONN4MST is completely different from other methods (**Fig. 1b**). While ONN4MST goes through the biome ontology to find the best possible source along different layers, other methods such as FEAST and SourceTracker treat all biomes as anarchically equal. The overall scheme of building the ONN model and using ONN4MST for source tracking is illustrated in **Supplementary Fig. 2**. Note that the contributions of every known biome would be estimated by the ONN model respectively.

### General model enables accurate source tracking with high scalability and stability

We constructed five datasets, representing sample collections with different numbers of biomes and samples, covering more than 100,000 real microbial community samples (**Supplementary Tables 1 and 2**). These five datasets contain samples from different niches including “Host_associated”, “Environmental” and “Engineered” as top biomes, which are representative of high-quality microbial community samples in public resources (**Supplementary Table 2**, **Methods**). Since these five datasets were designed to have varied complexities, each including different number of samples from different number of biomes, they could serve well for the evaluation of ONN4MST and other methods (**Fig. 2a**): The Combined dataset contains 125,823 samples and 114 biomes, which represents the largest datasets, as well as the largest model (the general model), used in this study. The FEAST dataset contains only 10,270 samples and 3 biomes. While the Human dataset, Water dataset, Soil datasets are respectively with moderate sample sizes (**Supplementary Tables 1**).

**Fig. 2:**
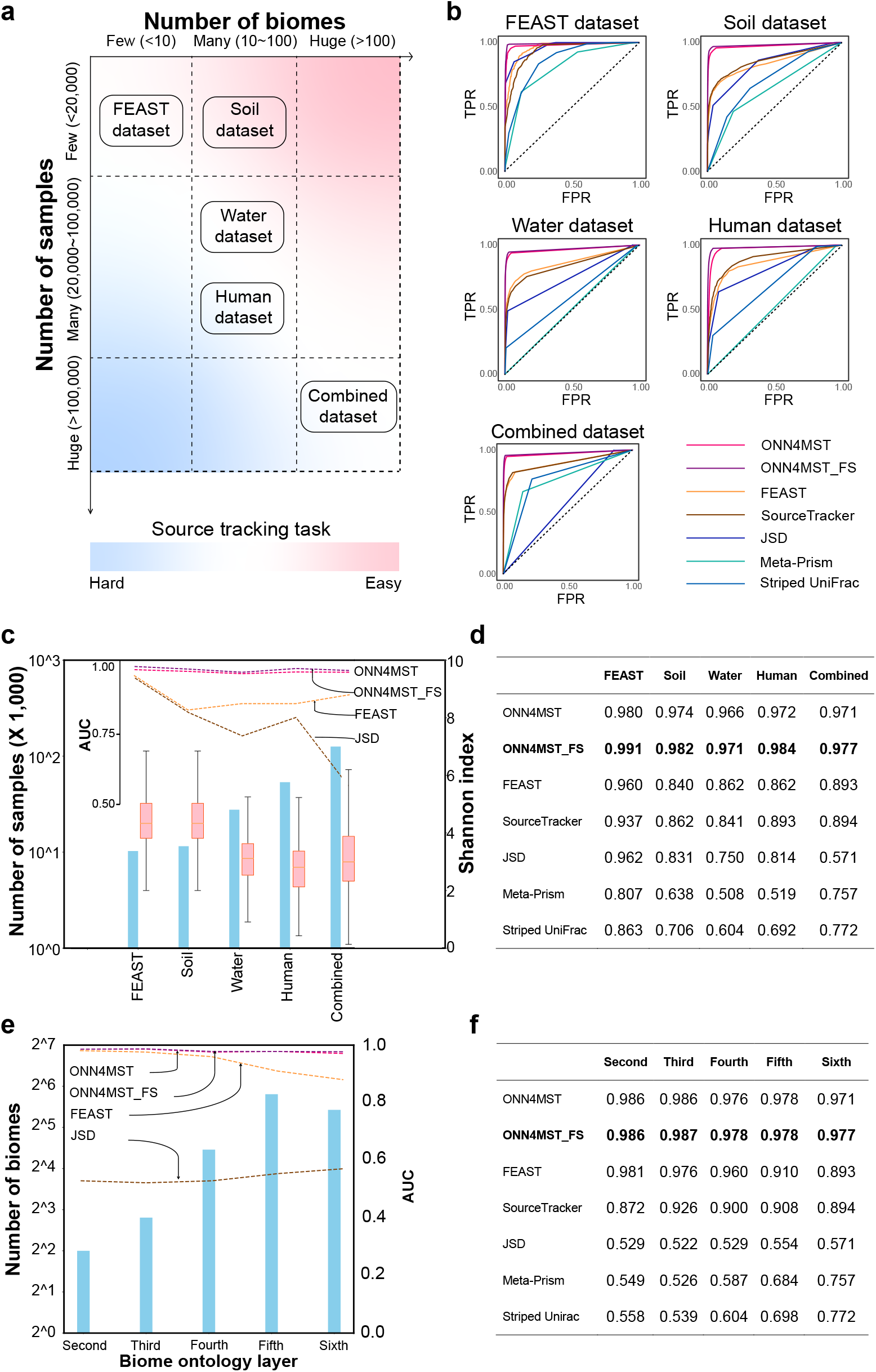
ONN4MST’s prediction accuracies are among the best on different datasets and different biome layers, while the performance of ONN4MST does not depend heavily on the number of biomes or number of samples in the dataset. **a.** The five datasets with varied complexities have provided source tracking tasks with different difficulties. **b.** The ROC curve of ONN4MST and other methods on all five datasets. **c.** The number of samples, the Shannon diversity and the source tracking results by different methods for the five datasets. The samples involved in each dataset are shown with blue bars, the Shannon diversity of each dataset is shown with red boxes, the AUC of several methods on each dataset is shown with dash lines. **d.** The AUC of all methods on all five datasets. **e.** The number of biomes and the source tracking results by different methods at different layers for the Combined dataset. The samples involved in each biome ontology layer are shown with blue bars, the AUC of different methods on each layer is shown with dash lines. **f.** The AUC of all methods at different layers. (**Abbreviations**. ONN4MST_FS: ONN4MST using selected features).

First and foremost, the performances of ONN4MST on all five datasets were evaluated. Results showed that the predicted biomes by ONN4MST were very close to the actual biomes, regardless of the datasets used for evaluation. For example, ONN4MST could achieve an accuracy of 0.99 and AUC of 0.97 on searching the Combined dataset with 125,823 samples from 114 biomes. When we applied ONN4MST on Human, Soil, Water and FEAST datasets, the accuracy and AUC of ONN4MST were also higher than 0.98 and 0.96 for these datasets (**Table 1**, **Supplementary Fig. 3**).

**Table 1.** Evaluation of ONN4MST on all five datasets. ONN4MST achieved the accuracy higher than 0.98 for all five datasets, and the AUC higher than 0.97 for all five datasets. **Note:** For each dataset, we used the model trained on that dataset for evaluation. The evaluation procedure of the ONN model is illustrated in **Supplementary Fig. 1c** and described in **Methods**. ONN4MST based on all features and selected features were both evaluated at the bottom (sixth) layer with a threshold of 0.5. (**Abbreviations**. Pr: Precision, Rc: Recall, Acc: Accuracy).

| Dataset | #Biomes | #Samples | All features |  |  |  |  | Selected features |  |  |  |  |
| --- | --- | --- | --- | --- | --- | --- | --- | --- | --- | --- | --- | --- |
| | | | Pr | Rc | Acc | $F_{max}$ | AUC | Pr | Rc | Acc | $F_{max}$ | AUC |
| Combined | 114 | 125,823 | 0.826 | 0.662 | 0.995 | 0.740 | 0.971 | 0.868 | 0.774 | 0.997 | 0.820 | 0.977 |
| Human | 25 | 53,553 | 0.822 | 0.521 | 0.984 | 0.695 | 0.972 | 0.894 | 0.826 | 0.991 | 0.863 | 0.984 |
| Water | 44 | 27,667 | 0.842 | 0.766 | 0.992 | 0.803 | 0.966 | 0.854 | 0.764 | 0.992 | 0.813 | 0.971 |
| Soil | 16 | 11,528 | 0.915 | 0.778 | 0.986 | 0.850 | 0.974 | 0.892 | 0.881 | 0.989 | 0.890 | 0.982 |
| FEAST | 3 | 10,270 | 0.793 | 0.795 | 0.984 | 0.803 | 0.980 | 0.895 | 0.812 | 0.989 | 0.862 | 0.991 |

ONN4MST based on selected features performed equally well or better than that based on all features. There are 44,668 taxa (or features) in total used in ONN4MST, while ONN4MST_FS (ONN4MST based on selected features) has utilized only 1,462 selected features (see **Methods** and **Supplementary Table 3**). Results showed that based on 1,462 selected features, ONN4MST_FS could attain slightly higher accuracy (0.997 vs. 0.995, on Combined dataset), AUC and *F_max_* compared to ONN4MST using all features (**Table 1**, **Supplementary Fig. 3**), which means that there is a certain degree of redundancy among all 44,668 features, and we can achieve the same accuracy with just 1,462 features compared with that using all 44,668 features. These results have emphasized the scalability and stability of the general model built based on the Combined dataset, either based on using all features, or using selected features.

Furthermore, we evaluated the universality of the general model built based on the Combined dataset, by applying it directly on the Human, Water, Soil, and FEAST datasets. It was found that the source tracking by using the general model was successful on those datasets which are composed of samples mostly from the Combined dataset’s samples (**Supplementary Table 4**, results on Human, Water, Soil datasets). However, when we applied the general model on datasets in which most of the samples were not previously observed in the general model or have more detailed biome ontology compared to the biome ontology used in general model, the general model would not perform well (**Supplementary Table 4**, results on FEAST dataset). Besides, results showed that it was unsuccessful when we applied the human model (the model built based on Human dataset) for source tracking on Soil and Water datasets (**Supplementary Table 5**).

### Comparison of ONN4MST and other source tracking methods

We then compared all six source tracking methods on all five datasets with different complexities (**Fig. 2a**). Results on all five datasets were evaluated seperately (**Fig. 2b,d**). Among the four datasets excluding FEAST dataset, ONN4MST was superior to other methods: ONN4MST reached an AUC of 0.97, while other methods only reached a maximum of 0.89 (**Fig. 2d**). As for the FEAST dataset, ONN4MST reached an AUC of 0.99, while other methods only reached a maximum of 0.96.

The performances of these methods on five datasets depend on the datasets’ complexities (**Fig. 2c**). While Soil dataset and Water dataset are among those with the highest Shannon diversity, the AUCs on these two datasets are also lower than those on Human dataset and Combined dataset. The high AUC on FEAST dataset is largely due to the small number of biomes used in FEAST dataset (**Supplementary Table 1**). On the other hand, the performance of ONN4MST on each dataset did not depend heavily on the number of samples in that dataset (provided that there are at least 10,000 samples in the dataset) (**Fig. 2c**, **Supplementary Table 1**). Furthermore, the prediction accuracies were not biased for certain biomes (provided that there are at least 100 samples in each biome) (**Supplementary Table 6**).

We further analyzed ONN4MST’s performances at different biome layers (**Fig. 2e,f**). Since it is the only method available that could source track samples at different layers of biome ontology, we have remolded other methods’ search scheme into a hierarchical prediction scheme (see **Methods**), so that their results are comparable to ONN4MST’s. Results have clearly shown that ONN4MST and ONN4MST_FS reached an AUC of 0.97 in minimum at all layers for the Combined dataset and these were noticeably superior to other methods (**Fig. 2e,f**). Thus, ONN4MST is not just the only method available that could source track the samples at different layers, but also the best method even when other methods were remoulded for such purpose.

### Running time and memory utilization benchmark

We evaluated the time and memory cost of all methods using a computational platform comprising Quadruplex E7-4809 v3 CPU with 315 GB RAM, Nvidia Tesla K80 GPU with 12 GB RAM. For time cost comparison, all actual times (search time, excluding I/O time) were converted to the equivalent time on a single core.

ONN4MST is superior to other methods in search time and memory utilization where the superiority expands as the number of source samples increases (**Fig. 3**). First of all, we tested the time cost by searching a single query against the five datasets respectively. For the Combined dataset including 125,823 source samples, ONN4MST and ONN4MST_FS took 0.18 seconds and 0.04 seconds, respectively, while distance-based methods took at least 1 second for a query. And FEAST took more than 100,000 seconds, and SourceTracker took even more time (**Fig. 3a**, on the Combined dataset, as also verified in Shenhav *et al*^9^.) Interestingly, though the time spent by FEAST and Source Tracker per thousand of source samples were both less than those reported in Shenhav *et al*^9^, these two methods costed magnitudes more time than ONN4MST (**Fig. 3a**). When we linearly extrapolated the number of source samples to one million in the dataset to be searched, the advantage of ONN4MST over other methods still held (**Fig. 3a**, hollow bars). When searching different number of queries against the Combined dataset, we observed the time cost follows this trend: supervised methods (ONN4MST and ONN4MST_FS) ≤ distance-based methods (JSD, Meta-Prism and Striped UniFrac) < unsupervised methods (FEAST and SourceTracker) (**Fig. 3b**). Again, when we linearly extrapolated the number of queries to one million in a batch, the advantage of ONN4MST over other methods still held (**Fig. 3b**, hollow bars).

**Fig. 3:**
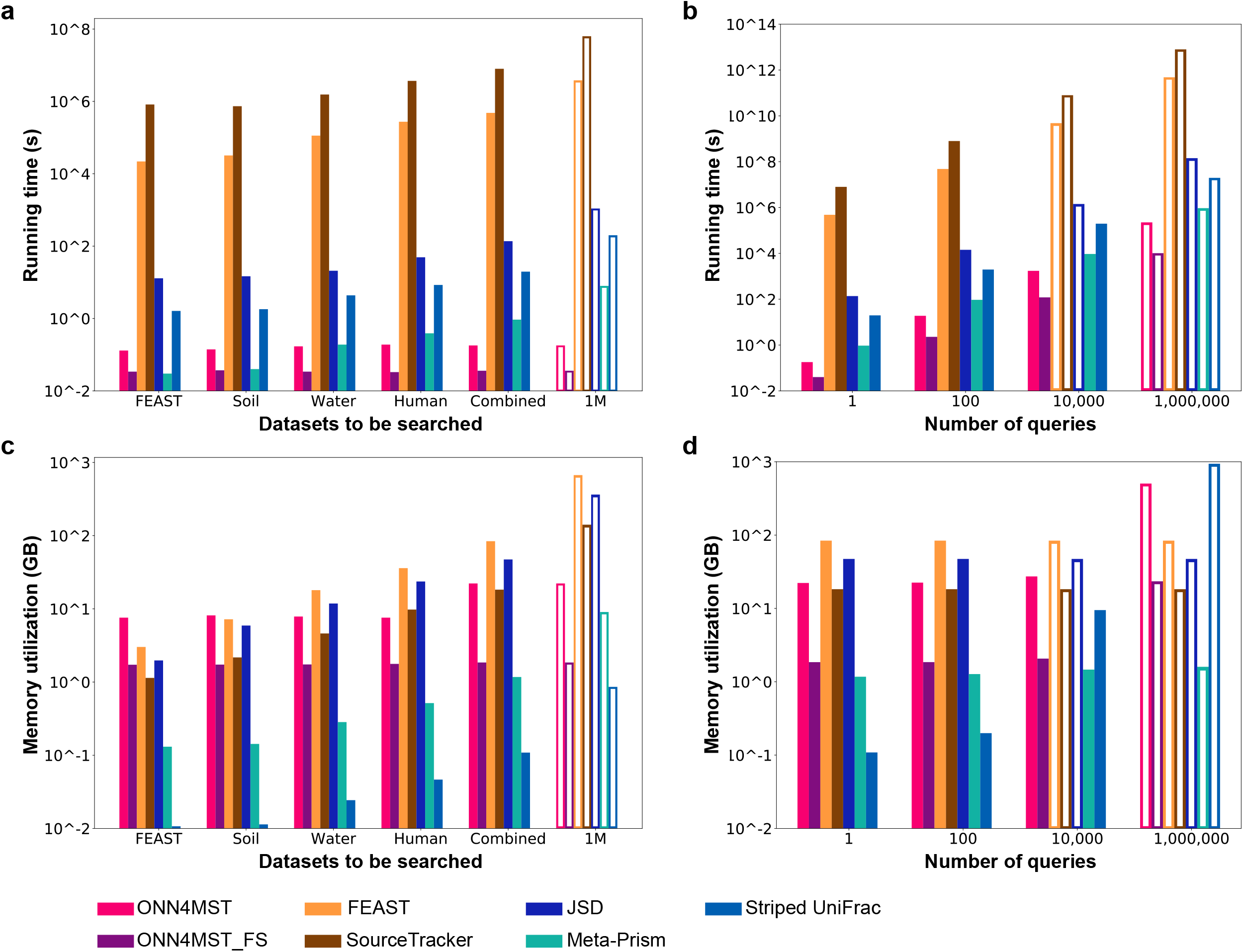
ONN4MST is superior to other methods in search time and memory utilization. **a.** Running time of different methods when search one query against different datasets. **b.** Running time of different methods when search queries of different sizes against Combined dataset. **c.** Memory utilization of all methods when search one query against different datasets. **d.** Memory utilization of all methods when search queries of different sizes against Combined dataset. **Note**: a hollow bar means that the value represent by this bar is the result of linearly extrapolation, both for running time and for memory utilization. (**Abbreviations**. ONN4MST_FS: ONN4MST using selected features, 1M: Results of linearly extrapolation with one million samples in use).

When memory utilization was evaluated, we have also observed the superiority of ONN4MST over most of the other methods. Specifically, when searching a single query against the Combined dataset, ONN4MST and ONN4MST_FS needed 22 GB and 2 GB of memory, respectively; while FEAST and SourceTracker needed 84 GB and 18 GB of memory, respectively; and JSD needed 47 GB of memory. Striped UniFrac and Meta-Prism (https://github.com/HUST-NingKang-Lab/Meta-Prism-2.0) were comparable with ONN4MST_FS in memory utilization, since they have optimized the data structure for sample comparison. When the number of queries in a batch exceeded 10,000, or the size of dataset to be searched varies, ONN4MST and ONN4MST_FS remain the ones that needed the least memory (**Fig. 3c,d**). Details about running time and memory utilization are presented in **Supplementary Tables 7-10**.

### Utility of ONN4MST in various source tracking applications

The objective of microbial community sample source tracking is knowledge discovery from the huge amount of microbial community samples of heterogeneous sources. Thus, we showcased the ability of ONN4MST in knowledge discovery from several perspectives: firstly, it can ensure accurate and interpretable source tracking, even on distinguishing samples from ontologically-close biomes; secondly, when samples’ biomes are previously less studied or unknown, ONN4MST could provide accurate clues for possible biome at higher layers, supplementing the information about such less-studies biome; thirdly, ONN4MST could help for accurate microbial contaminant detection; finally, “open search” of sample among the source samples with almost all possible biomes could identify similar samples from ontologically-remote biomes, leading to novel knowledge discovery.

### Centenarians share similar gut microbiota with young individuals

ONN4MST can distinguish samples from ontologically-close biomes, thus offers a quantitative way to characterize the development of human gut microbial community. In this context, we leveraged external sources of young individuals (30 years old on average) to understand the unique properties of gut microbiota in centenarians (persons over 100 years old). To demonstrate this capability, we first built a self-defined ONN model with two layers of biome ontology: “human gut” as first layer, while “Young human gut” and “Others or unknown” at second layer, through using a training set which contains 5,000 randomly selected human gut samples from the Combined dataset (**Supplementary Table 1**), together with 800 randomly selected human gut samples from young individuals in published studies^18,19^. Then, samples from centenarians (30 from Italy, and 51 from China)^18,19^ were used as queries for performing source tracking with the self-defined ONN model. Results revealed a significantly larger “Young human gut” contribution (Wilcoxon-test, p < 1e-3) in centenarians (**Supplementary Fig. 4**), regardless of the locations where these samples were collected, which were consistent with the results of published studies^18,19^. To prove that these gut microbiota properties were unique in centenarians, we have further collected 770 samples of normal seniors from another published study^20^ as queries for comparison. However, we could not observe the same phenomenon in these normal seniors (**Supplementary Fig. 4**).

Several other case studies that distinguish samples from ontologically-close biomes have also been conducted, with details in **Supplementary Note**, **Supplementary Figures 5 and 6**.

### Detecting microbial contamination in built environment

To validate ONN4MST’s ability on microbial contamination detection, we analyzed microbial community data collected by Lax *et al.*^21^ In this analysis, we investigated microbial contamination at several indoor house surfaces. We used skin samples from several body parts (skin, foot, hand and nose) and additional environmental, plants and mammal samples from the Combined dataset (**Supplementary Table 1**) as source samples, and samples from indoor house surfaces (“Bathroom Door Knob”, “Front Door Knob”, “Kitchen Counter”, “Kitchen Floor” and “Kitchen Light Switch”) as queries. Our analysis results by using ONN4MST have shown that microbial communities on these surfaces mostly originated from humans (**Fig. 4a**), largely in agreement with the original analyses of Lax *et al.^21^* using SourceTracker, and differs slightly from the results of Shenhav *et al^9^* These results were reasonable considering the strong influence of skin microbial communities on indoor house surfaces^22^, while they have again emphasized the challenge of source disambiguation for methods that do not consider ontology structure of the biomes. That is, treating each individual sample as an independent potential source would make differentiation of tiny sample differences among ontologically-close biomes impossible, thus underestimating the contributions of known sources at higher layers. We further investigated the composition of the unknown sources existed in **Fig. 4a**. In addition to the contribution of human, we found evidence for contributions from barley and bean product (0.6-1.1%) and marine product (0.2-0.4%) for kitchen environments, and potential evidence for contributions from agricultural (0.7-1.1%) and coastal (0.2-0.6%) for door knobs (**Fig. 4b,c**).

**Fig. 4:**
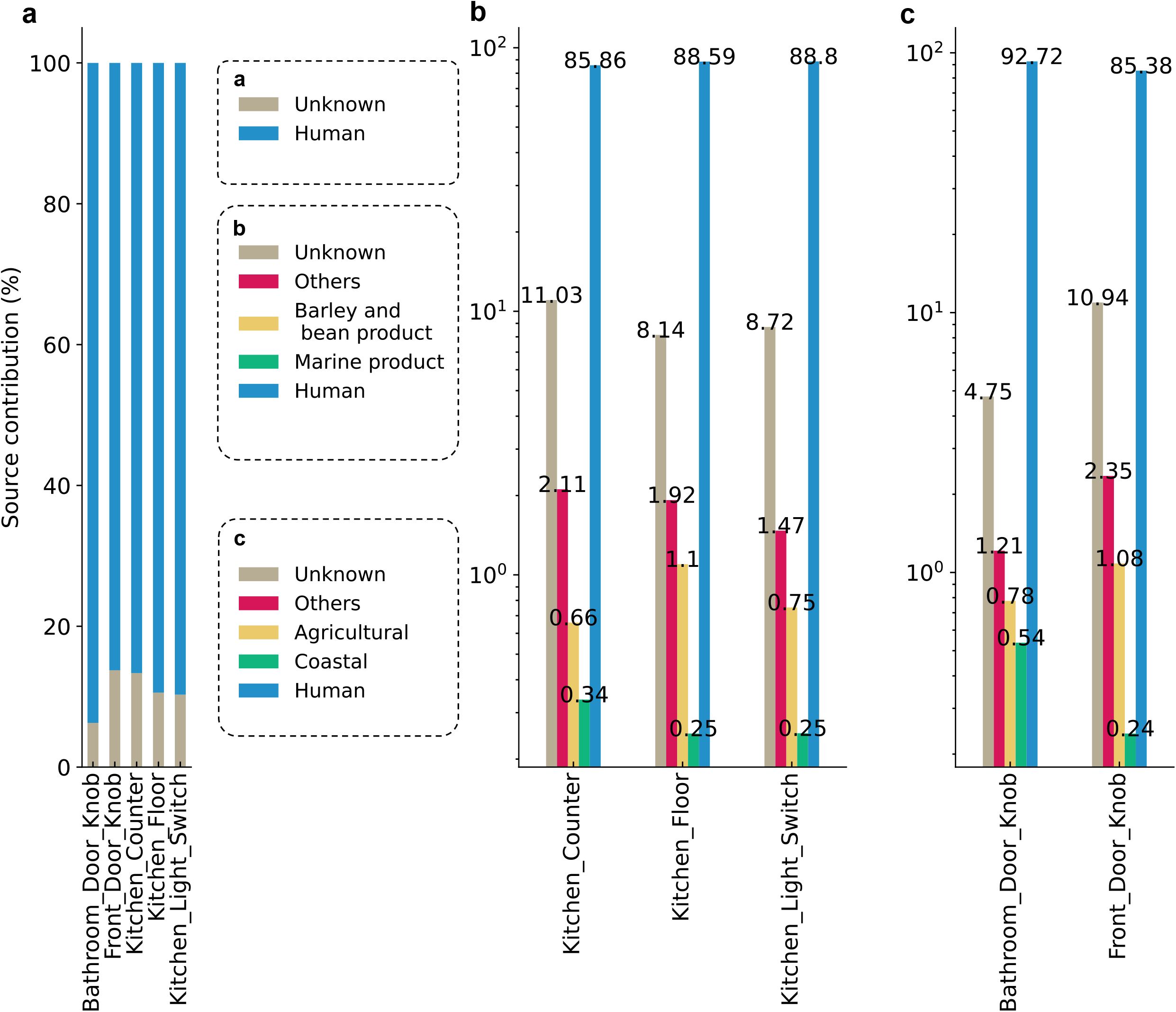
The contribution of the unknown sources in indoor house surface samples using ONN4MST. **a.** Mean source contributions considering 4 human skin sources (hand, foot, nose and skin-other across all inhabitants) using data from Lax *et al*.^21^ **b,c.** Further decomposition of the unknown sources existed in **Fig. 4a** has revealed other microbial con taminates in built environment.

### Source tracking of environmental samples from less studied biomes

This investigation was based on searching 11 groundwater samples from another published study^23^ (the biome “Groundwater” is less studied, with a handful of samples in the MGnify database, **Supplementary Table 2**) against the Combined dataset. ONN4MST could successfully identify the actual biome for the majority of these samples at different biome layers, such as “Aquatic” at the third layer and “Freshwater” at the fourth layer (**Fig. 5a-c**) (results at the fifth and sixth layers were shown in **Supplementary Fig. 7**). In contrast, FEAST and SourceTracker could not identify any source near “Groundwater”, while they only identified “Nutrient (Wastewater)” with the meaning marginally related with groundwater (**Fig. 5d,e**). Such differences in identification of actual biome are largely due to the fact that ONN4MST could screen the whole biome ontology, and identify possible sources at different layers, enabling it to at least identify the higher biome under which the actual biome belongs to, with high fidelity. Whereas FEAST and SourceTracker were designed without considering the biome ontology, they would assign “Unknown” for many of these samples. These results indicated that when the actual biome of sample was previously less studied, ONN4MST could provide accurate clues for possible biome at higher layers in the biome ontology, and such clues would become valuable assets in guiding the manual curation of these samples.

**Fig. 5:**
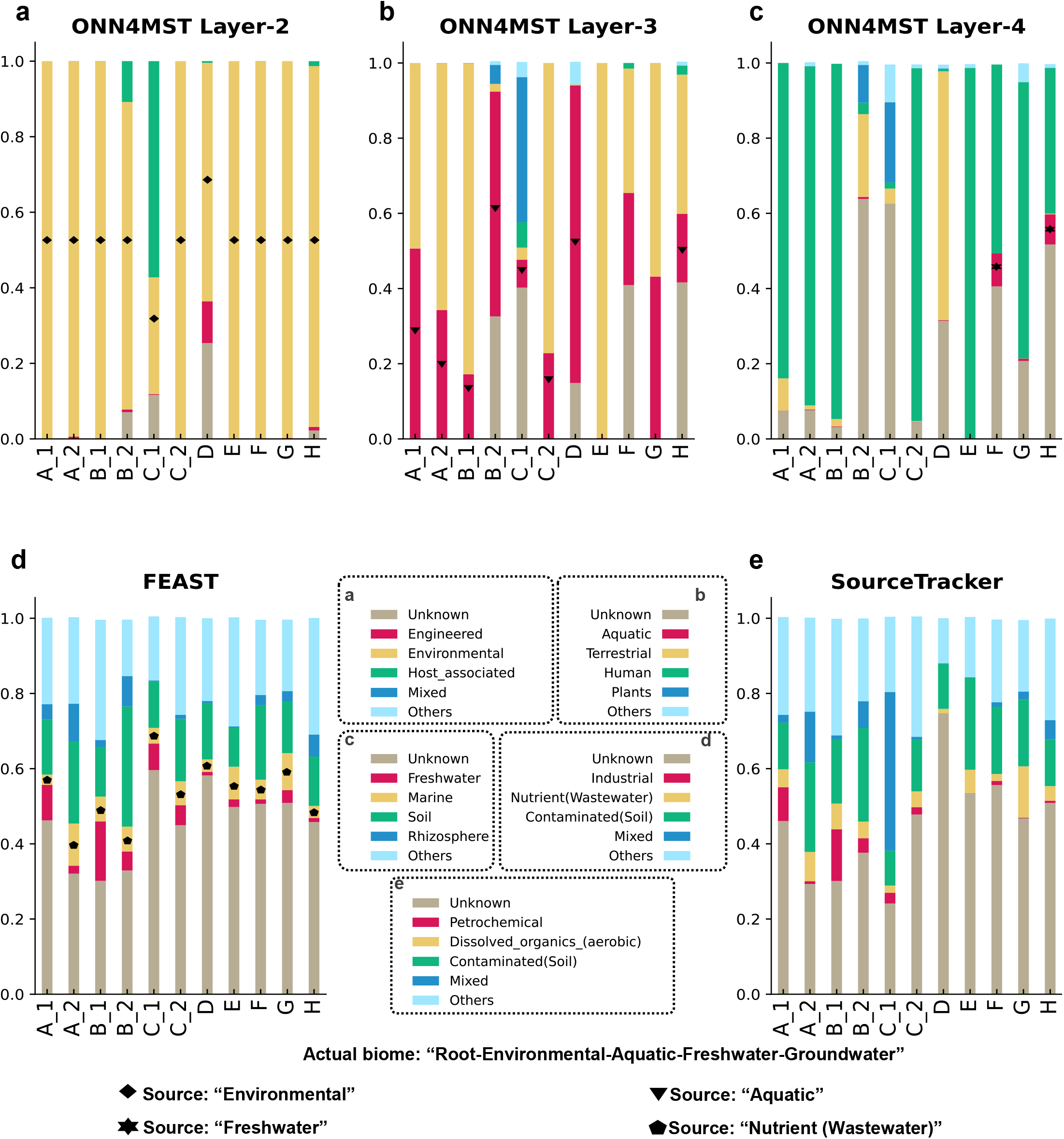
Successful source tracking of environmental samples from a less studied biome by using ONN4MST. Results were based on using 11 samples from groundwater environment, which represented a biome previously less studied. **a-c.** Source tracking results by using ONN4MST at the second, third and fourth layers; **d.** Source tracking results by using FEAST; **e.** Source tracking results by using SourceTracker. Actual biome of query sample: “Root-Environmental-Aquatic-Freshwater-Groundwater”. A_1, A_2: two samples collected from a single well; B_1, B_2: two samples collected from another single well; C_1, C_2: two samples collected from the third single well; D-H: samples collected from other five wells, respectively.

### Discovery of similar samples from ontologically-remote biomes

Another advantage of ONN4MST in source tracking is its ability for “open search” without any *a priori* knowledge about possible biomes where the query might be from, enabling it for novel knowledge discovery. We tested ONN4MST’s “open search” results, and found that it could discover similar samples among ontologically-remote biomes “Engineered”, “Host_associated” and “Environmental” (**Supplementary Table 11**). While some of the samples from the biome “Root-Environmental-Aquatic-Marine-Intertidal_zone” share similar environments (Baltic Sea) with the query sample from the biome “Root-Engineered-Wastewater-Industrial_wastewater-Petrochemi cal”, the literature has also verified that this query sample was marine-sourced “MGYS00005175” (from MGnify database). Such examples were plentiful (**Supplementary Fig. 8**), and many had very high contributions (> 0.8). However, there were also examples which might indicate possible mis-annotation or possible contaminations of samples in the MGnify database. For instance, more than 10 samples from the study “MGYS00001610” (from MGnify database) with annotated biome “Root-Engineered-Wastewater-Water_and_sludge” have been identified by ONN4MST as from biome “Root-Host_associated-Mammals-Digestive_system-Large_intestine-Fecal” (**Supplementary Fig. 8**). These results have verified our hypothesis that open search of sample among the source samples with almost all possible biomes could reveal remotely-similar samples, leading to novel knowledge that is never identified or interpreted before.

## Discussion

ONN4MST was designed to address the urgent need for fast, accurate and interpretable microbial community source tracking. It has been built based on an Ontology-aware Neural Network model, which has provided a solution for source tracking among sub-million samples and hundreds of biomes, outperforming state-of-the-art methods, thus enabling knowledge discovery from these heterogeneous samples. Microbial community sample source tracking has become increasingly important, largely due to the needs of source tracking in multiple areas. The requirements for high accuracy, high speed and high interpretability have thus become critical considerations for a successful source tracking method, especially when faced with the ever more complex situation where sub-million microbial community samples from hundreds of biomes are provided as possible sources for search.

The superiority of ONN4MST is established in several contexts. Firstly, ONN4MST is very robust against dataset heterogeneity: from a dataset with the number of biomes ranging from a handful to more than a hundred, as well as with the number of samples ranging from a few thousand to sub-million, it always provides the highest accuracies (AUC > 0.97) among state-of-the-art methods compared, making it the most scalable source tracking method. Secondly, based on the Human, Water and Soil datasets, the source tracking accuracies are all near-perfect (AUC > 0.97), indicating that ONN4MST could provide reliable insights for downstream analysis on implicating taxonomical or functional differences between healthy and diseased phenotypes, or on illuminating tiny differences among environmental samples from even slightly different niches. Furthermore, even when source tracking a sample against a database of sub-million samples, only less than 0.1 seconds is needed when we conduct ONN4MST search based on selected features, which is several orders of magnitude faster than other contemporary methods. Finally, the ability of ONN4MST for ‘open search’, without any *a priori* knowledge about possible biomes where the query might be from, enables it for interpretable knowledge discovery.

The advantage of ONN4MST over other state-of-the-art source tracking methods is essentially dependent on two technical advancements: the deep learning model, and the ontology structure. Though the currently ongoing shift towards supervised learning methods is not surprising for the source tracking research, the superior performance of ONN4MST over existing methods is still quite pronounced. ONN4MST’s advantage also stems from its consideration of the ontology structure of the biomes: by embedding the ontology considerations into the ONN learning model, ONN4MST naturally becomes suitable for solving the ontology relationships among biomes.

ONN4MST is not without limitations. Most importantly, the accuracy of ONN4MST is heavily dependent on the ONN model built based on existing biome ontology information. If there comes a new biome ontology with more detailed biomes involved (for example, if we need to refine the source tracking results to human gut down, to differentiate niches such as adult’s gut from infant’s gut), or simply with more biome relationships involved, then the ONN model should be re-trained for accurate source tracking. Such biome ontology-wide scalability problem could potentially be solved by Transfer Learning approaches.

In summary, ONN4MST is an ontology-aware deep learning method that has pushed the envelope of microbial source tracking, enabling near-optimal accurate, ultrafast and interpretable source tracking. ONN4MST has enabled in-depth pattern and function discoveries among sub-million microbial community samples, allowing for tracking the potential origin of microbial community with diverse niche background, as well as distinguishing samples from different health conditions or diverse environments. Thus, it could have a broader area of application, such as contamination screening, novel or refined biome discovery, new functional microbiome discovery, and even source tracking of biomes from which protein sequences could be supplemented for computational protein 3D structure prediction^24,25^.

## Methods

### Datasets

We evaluated the performances of ONN4MST and other source tracking methods based on five different datasets (**Supplementary Table 1**). These five datasets comprise samples from different niches, which are representative of high-quality samples in public resources.

The “Combined dataset” consists of 125,823 microbial community samples collected from EBI MGnify database (https://www.ebi.ac.uk/metagenomics/), accessed as of January 2020 (**Supplementary Table 1**). This is a comprehensive dataset containing samples from 114 biomes (**Supplementary Table 2**), and the 125,823 microbial community samples represent more than half of the samples in EBI MGnify (as of January 1st, 2020). These samples contain taxonomical information for 225 phyla, 6,232 families, 16,081 genera and 45,477 species.

The “Human dataset” consists of 53,553 microbial community samples selected from the Combined dataset, representing a subset of samples from the human niches (**Supplementary Table 1**). Specifically, these samples are collected under these biomes: “Root-Host_associated-Human-Skin”, “Root-Host_associated-Human-Circulatory_system”, “Root-Host_associated-Human-Digestive_system” and “Root-Host_associated-Human-Reproductive_system” (biomes at higher layer). This dataset contains 53,553 samples from a total of 25 biomes. These samples contain taxonomical information for 204 phyla, 2,801 families, 6,523 genera and 16,135 species.

The “Water dataset” consists of 27,667 microbial community samples selected from the Combined dataset, representing a subset of samples from the water niches (**Supplementary Table 1**). Specifically, these samples are collected under these biomes: “Root-Environmental-Aquatic-Freshwater”, “Root-Environmental-Aquatic-Marine” and “Root-Environmental-Aquatic-Non-marine_Saline_and_Alkaline” (biomes at higher layer). This dataset contains 27,667 samples from a total of 44 biomes. These samples contain taxonomical information for 222 phyla, 6,040 families, 15,261 genera and 36,406 species.

The “Soil dataset” consists of 11,528 microbial community samples selected from the Combined dataset, representing a subset of samples from the soil niches (**Supplementary Table 1**). Specifically, these samples are collected under these biomes: “Root-Environmental-Terrestrial-Soil”, and “Root-Host_associated-Plants-Rhizosphere” (biomes at higher layer). This dataset contains 11,528 samples from a total of 16 biomes. These samples contain taxonomical information for 201 phyla, 2,962 families, 6,753 genera and 12,769 species.

These three datasets (Human, Water and Soil datasets) were designed with several reasons in consideration. Firstly, these three datasets are representative enough and frequently-used subsets^11^ from the Combined dataset. Secondly, these three datasets are also distinct, since the Alpha diversity of samples from each of these datasets is significantly different from the other two: while samples from soil niches are considered more complicated, those from human and water niches are considered less so. Finally, samples from these niches are more comprehensively explored than other less studied niches, and they are of relatively higher quality of samples from these three niches.

The “FEAST dataset” consists of 10,270 microbial community samples selected from the datasets used in the Lax *et al*.^9^ (**Supplementary Table 1**). Specifically, these samples are all collected from three biomes (“Root-Host_associated-Human”, “Root-Host_associated-Human-Digestive_system-Large_intestine-Fecal” and “Root-Mixed”). These samples contain taxonomical information for 133 phyla, 1,118 families, 3,389 genera and 5,762 species. The “FEAST dataset” is the smallest dataset used in this study, and it is the simplest dataset with regard to the number of biomes involved. Yet it is a dataset of unique importance, as the source tracking methods evaluated in this study could be benchmarked on this medium-sized and credible human gut dataset^9,15^ for fair assessment of accuracy and efficiency.

### Data representation

we generated the Matrix for each microbial community sample, so that the abundances for all taxa at seven taxonomical levels including super-kingdom, kingdom, phylum, class, order, family, and genus (simply referred to as “sk”, “k”, “p”, “c”, “o”, “f”, and “g”) can be retained. The abundance of taxa at different levels were filled in the Matrix (**Figure 1**). Within the Matrix, seven columns respectively represent seven taxonomical levels. And 44,668 rows respectively represent relative abundance for 44,668 taxa (also referred to as features). For a detailed description and an example of the data representation, see **Supplementary Note** and **Supplementary Table 3**.

### Feature selection

To improve the efficiency and accuracy of ONN4MST, we conducted feature selection by using a random forest regression model (Python-3.7.4 and Scikit-learn-0.22.1). An abundance-based pre-filtering and an importance-based selection were performed in sequential order. In doing so, we treated each row (representing the abundances of a taxon, see **Supplementary Table 3**) of the Matrix as a feature. Then, a series of adaptive thresholds (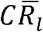 and 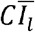) were applied to different taxon levels, in which 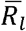 and 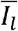 stand respectively for the relative abundance and the feature importance. *level* ∈ {*sk, k, p, c, o, f, g*} and the coefficient *C* was set to 0.001. As a result, we have selected 1,462 features with relative abundance and feature importance above the thresholds from all 44,668 features involved in this study.

### Biome ontology

We constructed a comprehensive biome ontology using 114 biomes (**Supplementary Table 2**) collected from EBI MGnify database (https://www.ebi.ac.uk/metagenomics/biomes). In this process, we organized the biome ontology as a tree, by treating a biome with multiple parent biomes in the higher layer (e.g. “Human-Digestive_system” and “Mammal-Digestive_system”) as seperate biomes. Next, the ontology tree containing 6 layers and 133 nodes (representing 114 biomes) was constructed, by using Python-3.7.4 and Treelib-1.5.5. As a result, each biome was represented by at least one node in the ontology tree. The ontology tree has “Root” at the first layer, biomes (nodes) including “Environmental”, “Host_associated”, and “Engineered” at the second layer, and 7, 22, and 56 biomes (nodes) at the third to fifth layers respectively, with 43 biomes (nodes) including “Coral reef’, “Fecal” and “Saliva” at the bottom (sixth) layer (**Supplementary Table 2).**

### Sample Labeling

In all experiments, we used microbial samples each with a label annotated by using 6-layers biome ontology to validate our model. For example, there are 22 samples labeled as “Root-Host_associated-Human-Digestive_system-Oral-Throat” in the Combined dataset (by separating different layers with the “-” symbol).

### Building ONN model

We used Tensorflow-1.14^26^ to build and train our Ontology-aware Neural Network model. Our model was trained on a computational platform comprising Quadruplex E7-4809 v3 CPU with 315 GB RAM and Nvidia Tesla K80 GPU with 12 GB RAM.

Ontology-aware Neural Network has four conceptual modules in total: a feature extraction module for basic feature extraction, a feature encoding module for layer-specific feature encoding, a feature integration module for inter-layer information integration, and an ontology prediction module for ontology walk through and source contribution calculation (**Supplementary Fig. 1a**). The feature extraction module accepts a sample represented by the Matrix, extracts the feature information from the Matrix and deliver them to the feature encoding module. The feature encoding module consists of a series of fully-connected layers. It accepts the output of feature extraction module, and encodes layer-specific feature information for each of the six biome ontology layers. The feature integration module consists of several fully-connected layers, which serves for inter-layer information integration. The ontology prediction module consists of five sigmoid layers (corresponding to the 2^nd^, 3^rd^, 4^th^, 5^th^ and 6^th^ biome ontology layers), each sigmoid layer accepts the output of feature encoding module and computes the contribution of all biome sources on its corresponding biome ontology layer.

We chose 8-fold cross validation for model training and testing (**Supplementary Fig. 1c**). For each dataset, we randomly split it into 8 folds, each fold including a training set (87.5%) and a testing set (12.5%). For each fold, the model was trained (in batches of 512 samples) for 30,000 iterations or until training accuracy converged, and the model with the highest accuracy on the training set was selected for testing. The results on the testing set are organized in the form of a hierarchical prediction (with prediction results from 2^nd^ to 6^th^ layers), which would then be evaluated.

### Other methods used in this study

Three distance-based methods: JSD, Striped UniFrac and Meta-Prism, two unsupervised machine learning methods: Expected-Maximization based method FEAST and Bayesian based method SourceTracker; as well as our supervised deep learning method (ONN4MST), were applied for microbial source tracking. In this study, the source tracking results (predicted biomes) of multiple methods were compared against the microbial community samples’ actual source (actual biomes).

The distance-based methods are based on pair-wise calculation of sample distances, and such methods depend heavily on the presence of species and their relative abundance for individual samples, regardless of weighted or unweighted scoring functions used. Among distance-based methods, JSD does not consider the phylogenetic relationships among species, while methods such as Striped UniFrac and Meta-Prism do (we have used Meta-Prism 2.0 for comparison in this study). However, distance-based methods have a binomial increase in time cost with the increase of the number of samples.

Unsupervised methods for microbial community sample comparison are based on profile-based statistical models, either the Bayesian model used in the SourceTracker method, or the Expected-Maximization (EM) model used in the FEAST method. Unsupervised methods are typically more accurate than distance-based methods. However, since unsupervised methods still do not consider the intricate but important patterns of a set of samples from similar niches, their tolerance to noisy signals in samples is not high, hence potentially would lead to biased mismatches. Details about the source tracking methods other than ONN4MST used in this study are provided in **Supplementary Note**.

### Hierarchical prediction

In order to carry out comparison of ONN4MST against other methods at different layers of biome ontology, all other methods were remolded, so that the prediction results of these methods (excluding ONN4MST) at different layers could be produced. Based on the source contributions of biomes at the sixth (bottom) layer, the source contributions of biomes for other layers were computed using *P_f_* = Σ_*f_c_∈c_f_*_*P_f_c__*. Where *P_f_* is a source contribution for *f, C_f_* is a set of children biomes for biome source *f* in the biome ontology. *f_c_* is a child biome of *f*. We used NumPy-1.18.1 and Treelib-1.5.5 in the process.

### Benchmarking measures

To benchmark and compare the results based on ONN4MST and the other five methods, we used these measures:

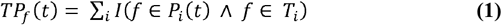

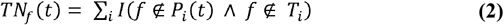

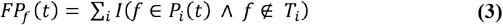

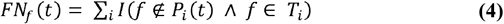

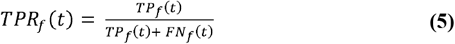

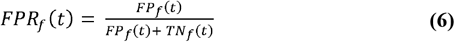

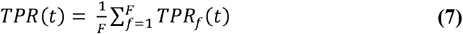

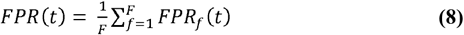

where *f* is a biome source, *P_i_*(*t*) is a set of predicted biomes for a microbial community sample *i* and threshold *t* ∈ [0,1] with a step size of 0.01, *T_i_* is a set of actual biomes for a sample *i, F* is the total number of biomes, and *I* is a logical operation function, the value of *I* is 1 when the result of logical operation is TRUE, else 0.

Four evaluation metrics *(Accuracy, Precision, Recall* and *F_max_*) were introduced. These evaluation metrics are computed with the following formulas:

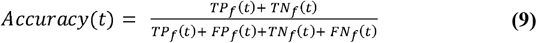

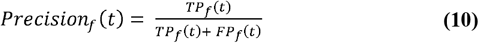

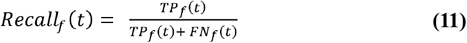

where *TP* is true positive, *TN* is true negative, *FP* is false positive, *FN* is false negative. Subsequently, we compute Fl for threshold *t* ∈ [0,1] with a step size of 0.01 by using the average precision and average recall for all actual biomes that we predicted at least one time. Then, we select the maximum *F*l as *F_max_*. These evaluation metrics are computed with the following formulas:

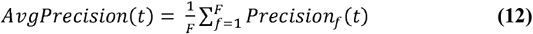

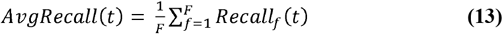

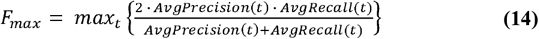

Then, ROC (Receiver Operating Characteristic) curves, which are based on contrasting the true positive rate (TPR) against the false positive rate (FPR), were plotted. AUC (Area Under the Curve) reflects the ability of model to correctly predict the biomes (sources) of microbial community samples. AUC is calculated with the following formula:

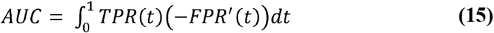

## Data availability

The selected samples from Combined dataset, which were assigned to Human dataset, Water dataset, Soil dataset respectively, were annotated with their respective assignments in **Supplementary Table 2**. Data download links are provided in **Supplementary Table 12**.

## Code availability

All source codes have been uploaded to the website at: https://github.com/HUST-NingKang-Lab/ONN4MST. Detailed parameters of software and package we used in this study are provided in **Supplementary Table 13**.

## Acknowledgments

We are grateful to Chuanle Xiao, Jianyang Zeng and Qingyang Yu for insightful discussions. This work was partially supported by National Science Foundation of China grant 81774008, 81573702, 31871334 and 31671374, and the Ministry of Science and Technology’s national key research and development program grant (No. 2018YFC0910502).

## Author contributions

KN conceived of and proposed the idea, and designed the study. YGZ, HC, HQ, KK, YZD, ZXC performed the experiments and analyzed the data. YGZ, HC, KN and XC contributed to editing and proof-reading the manuscript. All authors read and approved the final manuscript.

## Ethics approval and consent to participate

Not applicable

---

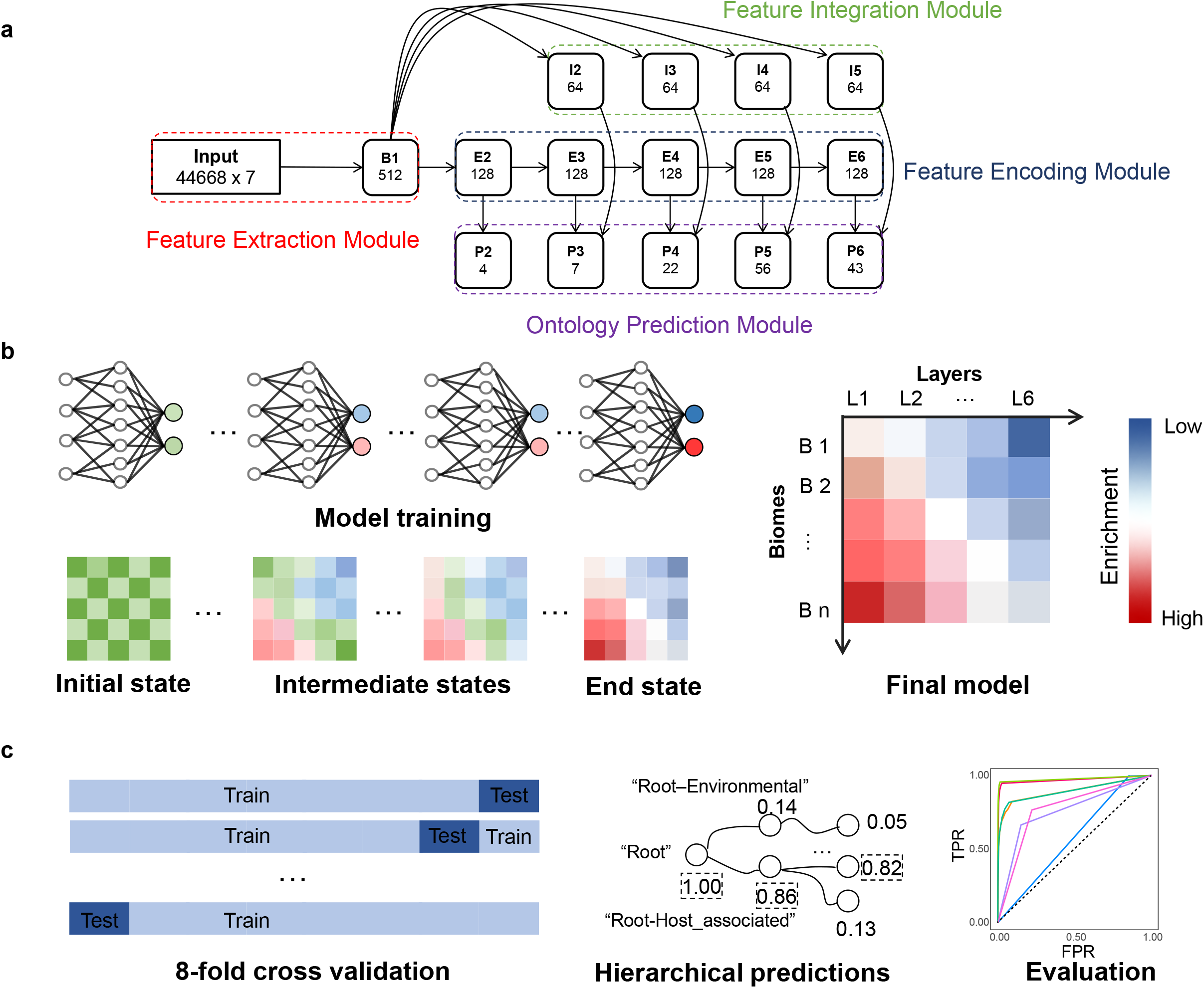

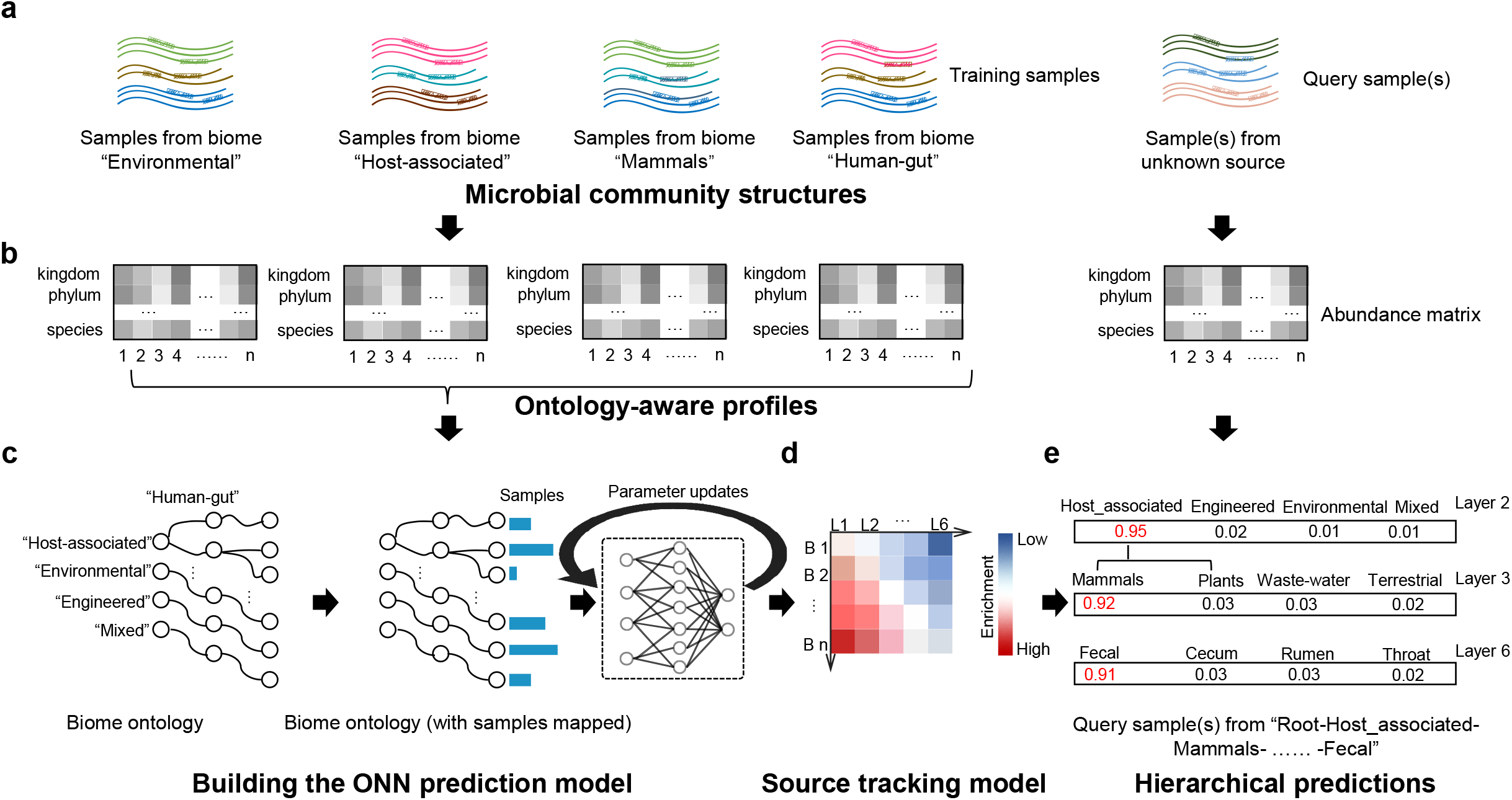

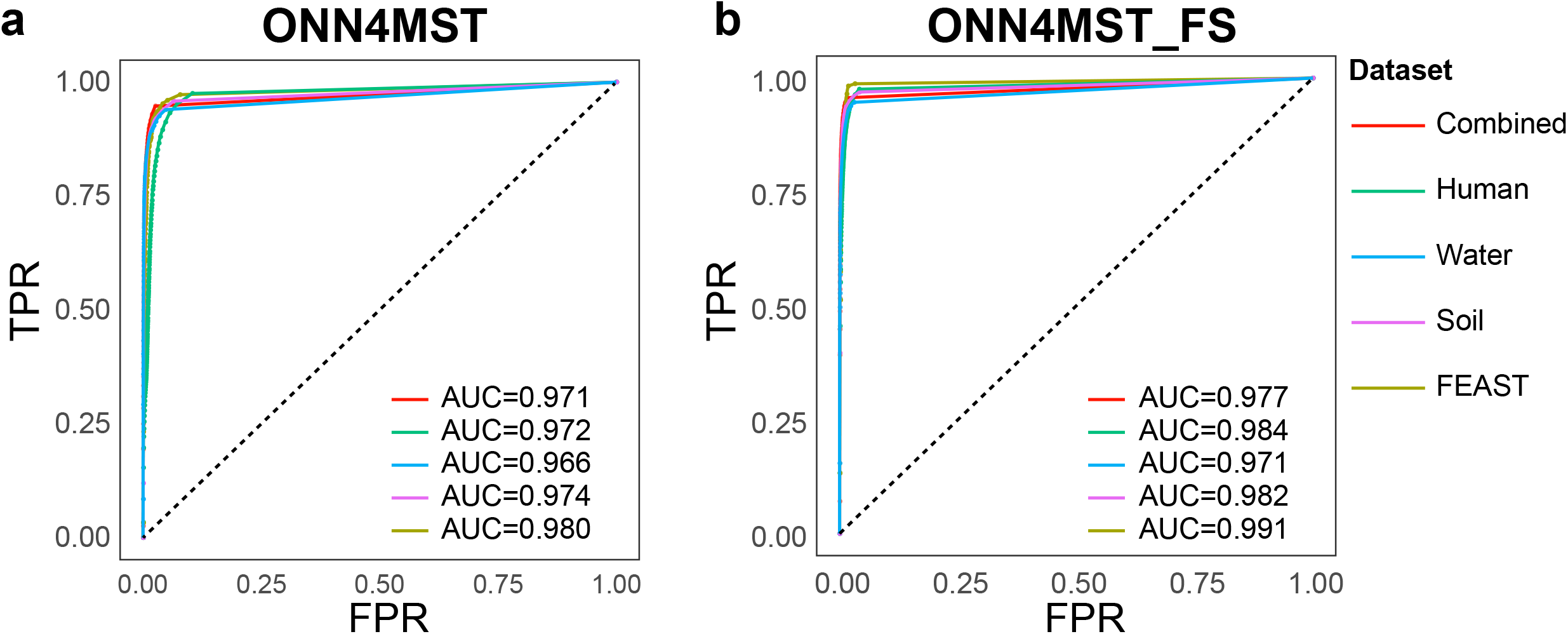

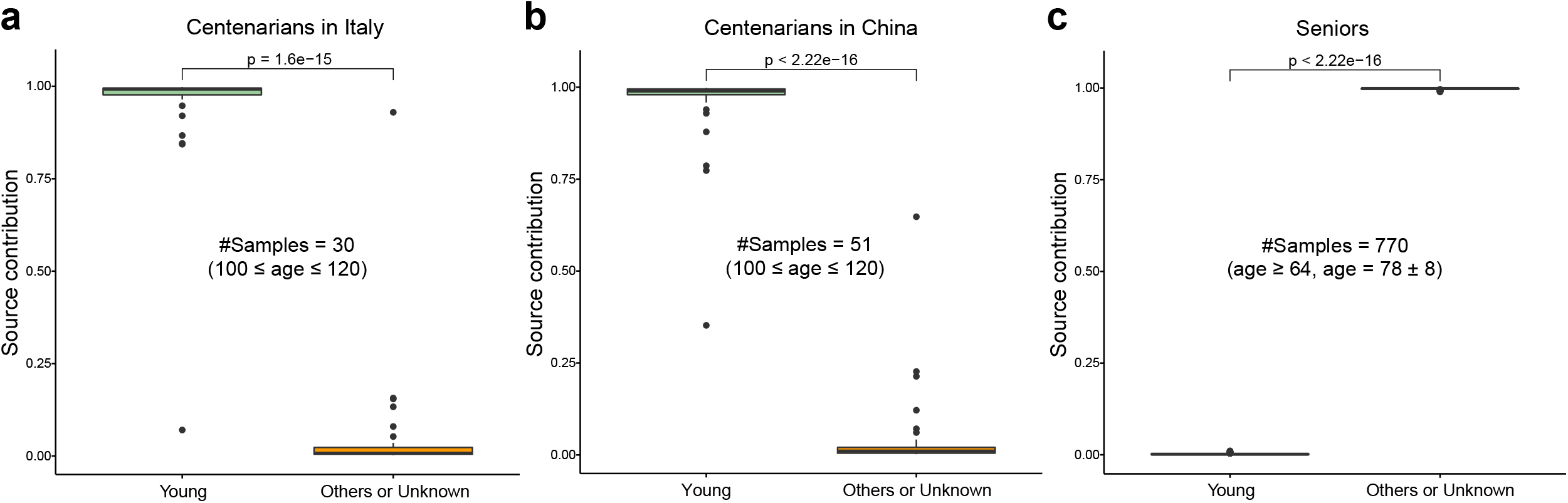

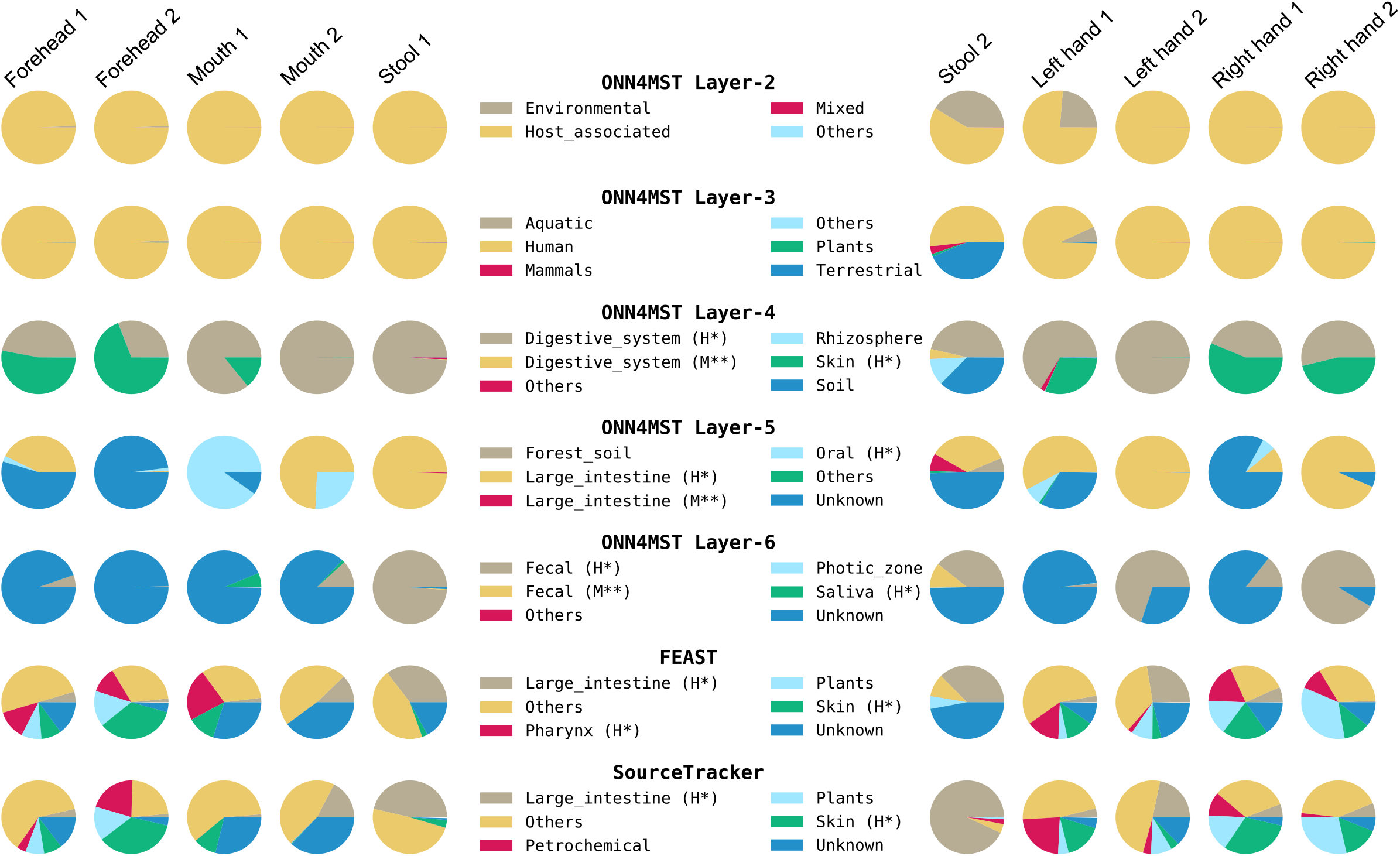

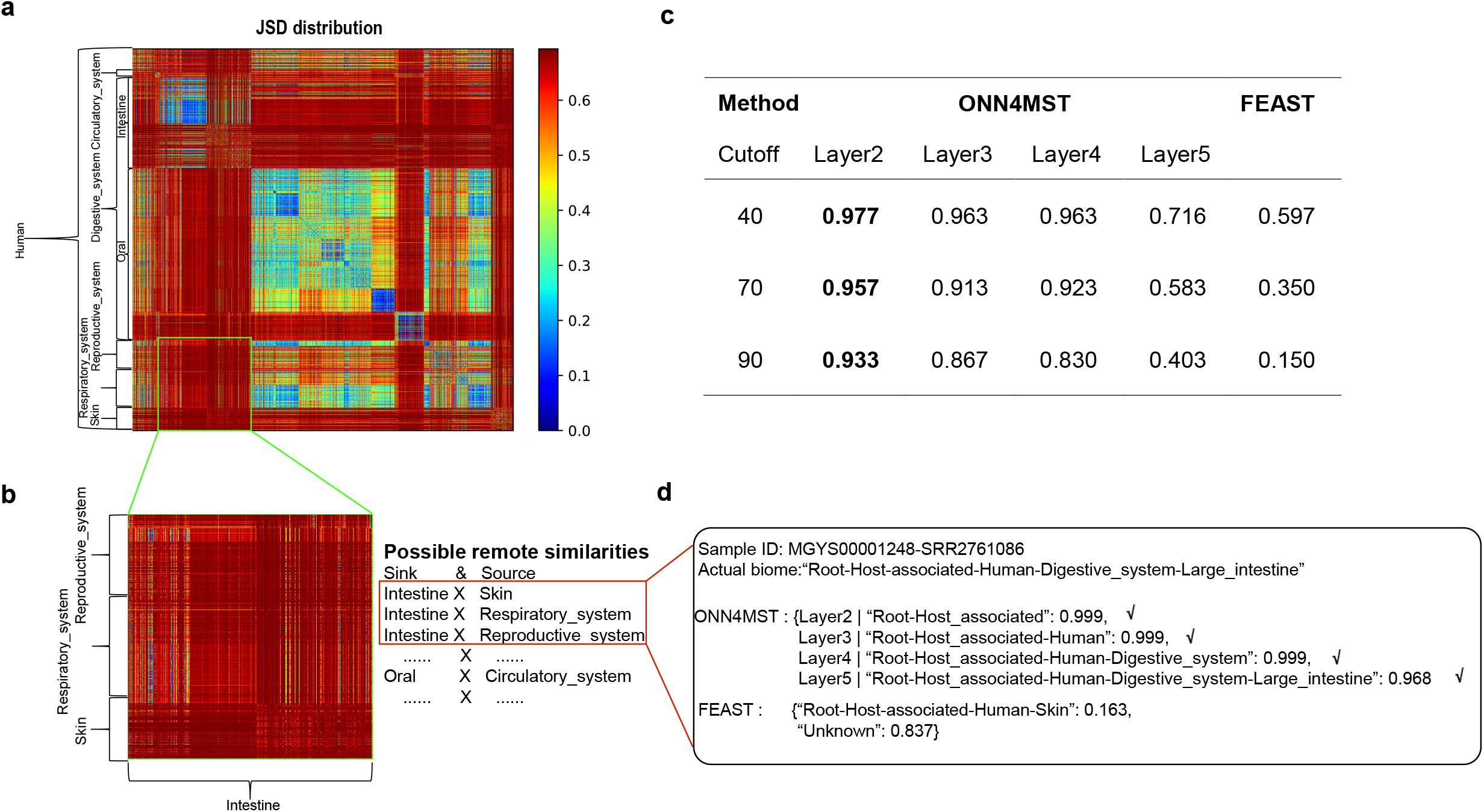

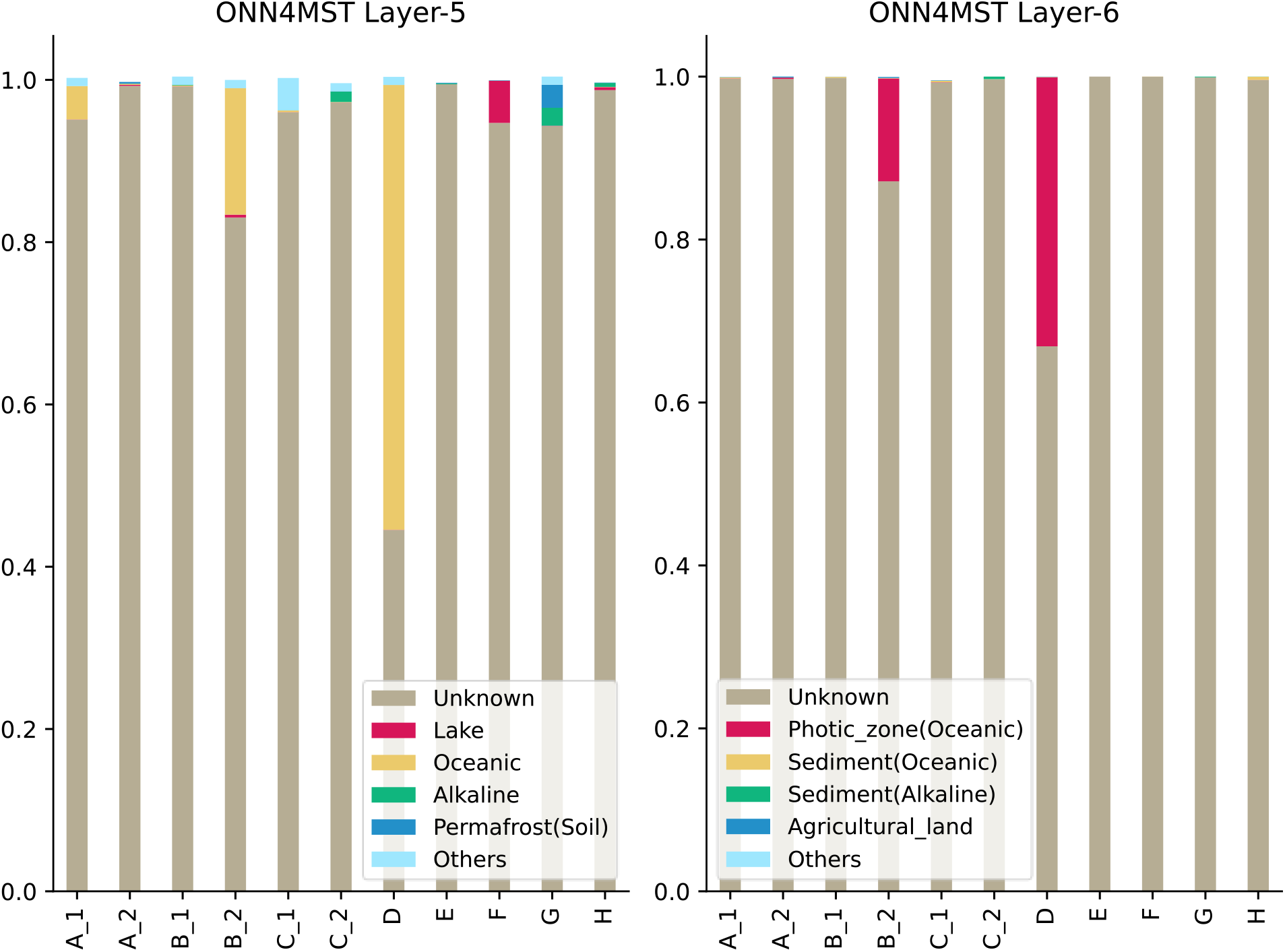

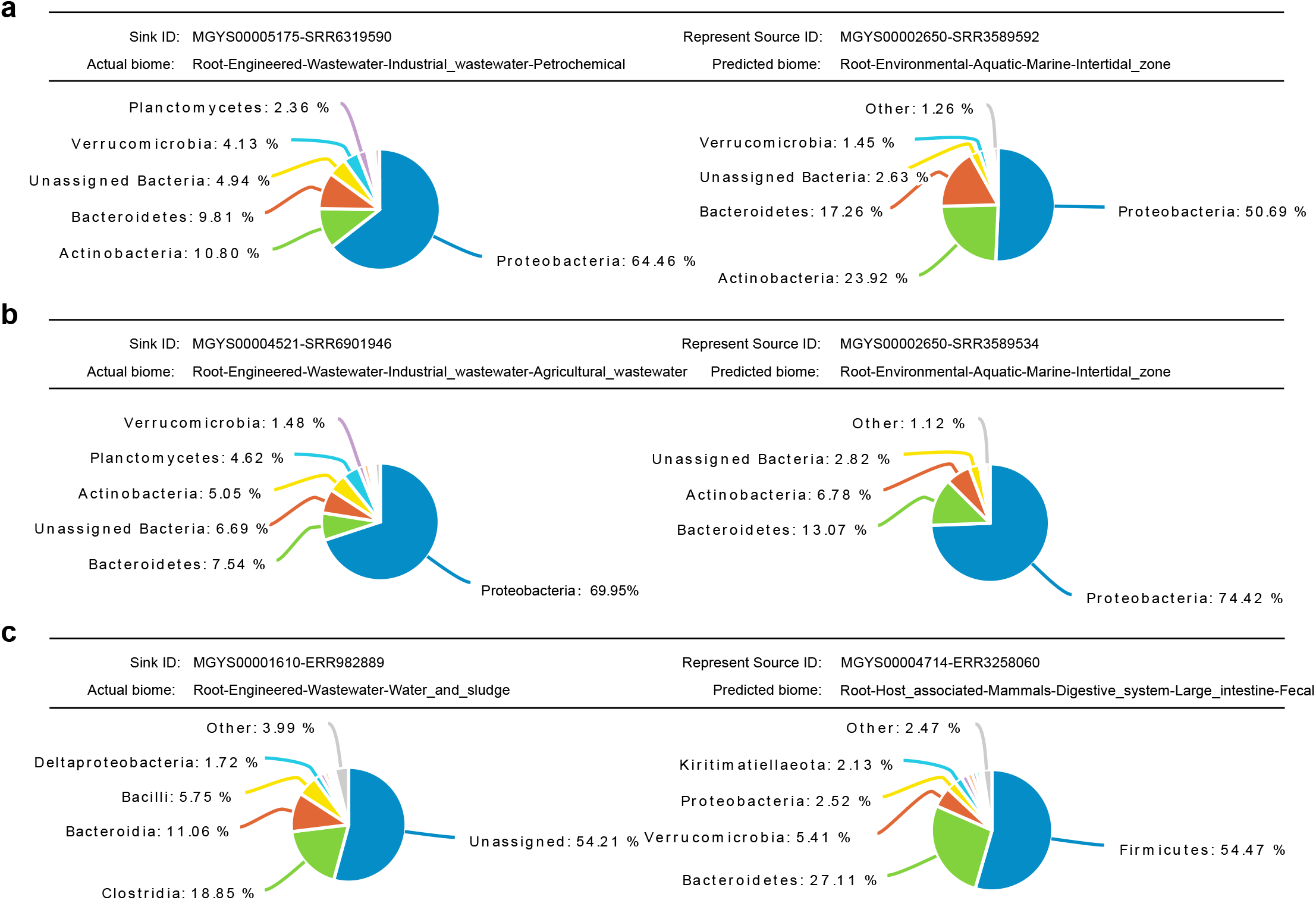

