## Supplementary Information for "Ontology-Aware Deep Learning Enables Ultrafast, Accurate and Interpretable Source Tracking among Sub-Million Microbial Community Samples from Hundreds of Niches"

Supplementary Figures 1-8, Supplementary Tables 1-13 and Supplementary Notes 1-3

**Supplementary Figure 1**

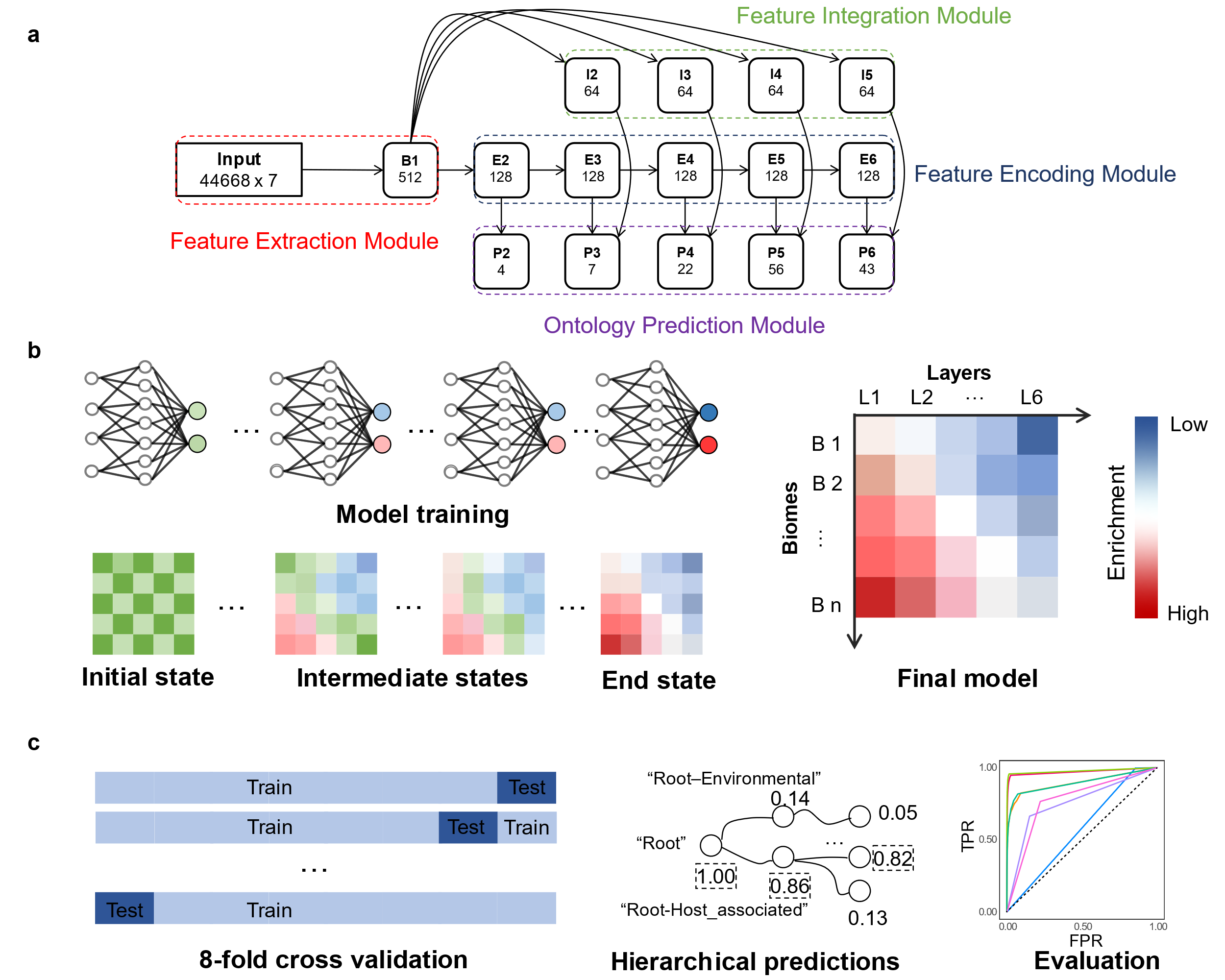

**Supplementary Figure 1.** **The conceptual modules, the training procedure, and the evaluation procedure of the ONN model.** **a.** The conceptual modules of the ONN model. Feature Extraction Module is designed for basic feature extraction. Feature Encoding Module is designed for layer-specific feature encoding. Feature Integration Module is designed for inter-layer information integration. Ontology Prediction Module is designed for ontology walk through and source contribution calculation. **Note:** Each node is annotated with a node ID (top half) and its data dimension (bottom half). In feature extraction module, the dimension of the input (the Matrix) is 44,668×7, representing 44,668 taxa (at different taxonomical levels) and 7 taxonomical levels. B1: the node serving for basic feature extraction, represented by 512 features. In feature integration module, each node (I2, I3, I4 and I5) corresponds to a biome layer-layer connection (2^nd^-3^rd^, 3^rd^-4^th^, 4^th^-5^th^ and 5^th^-6^th^, respectively), represented by 64 features. In feature encoding module, each node (E2, E3, E4, E5 and E6) corresponds to a specific biome layer (2^nd^, 3^rd^, 4^th^, 5^th^, and 6th, respectively) information, represented by 128 features. In ontology prediction module, each node represents a prediction results at one biome layer, represented by the number of biomes for that layer. For example, there are 4, 7, 22, 56, 43 biomes (nodes) in the 2^nd^, 3^rd^, 4^th^, 5^th^ and 6^th^ layer in the biome ontology, and each biome would have a prediction result accordingly. **Note:** we organized the biome ontology as a tree, by treating a biome with multiple parent biomes in the higher layer (e.g. “Human-Digestive_system” and “Mammal-Digestive_system”) as seperate biomes. **b.** The training procedure of the ONN model. From left to right: The ONN model was randomly initialized at the initial state. Then, the ONN model was trained for 30,000 iterations, and the model in these iterations that reached the highest accuracy on the training set was selected as the final model. **c.** The evaluation procedure of the ONN model. Left: The dataset was randomly split into 8 folds, each fold including a training set (87.5%) and a testing set (12.5%). Middle: For each fold, the model was trained on the training set, and the testing results on the testing set are organized in the form of a hierarchical prediction. Right: These hierarchical prediction results would then be evaluated.

**Supplementary Figure 2**

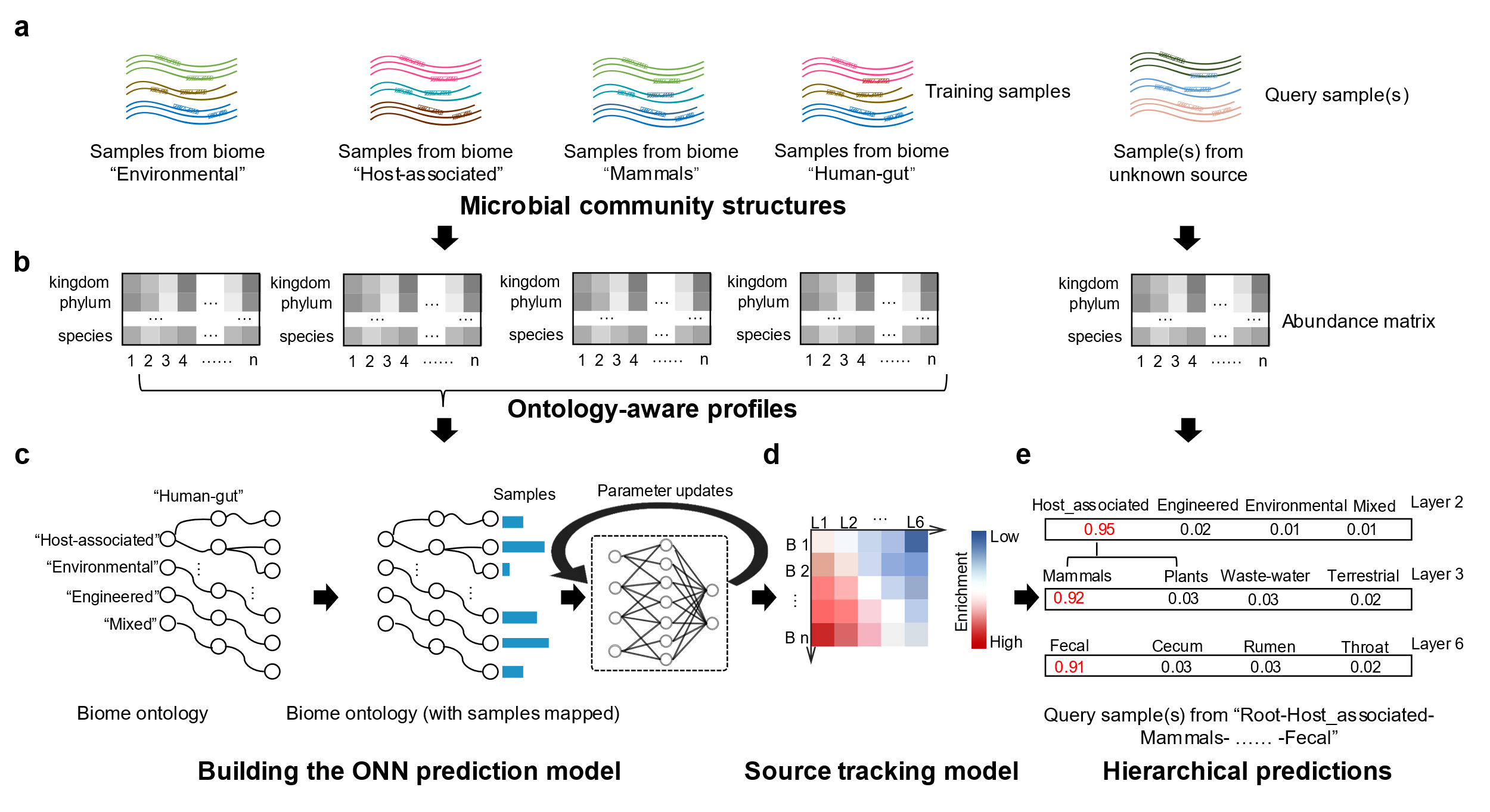

**Supplementary Figure 2. Overview of using ONN4MST for microbiome sample source tracking. a.** Collecting microbial community samples; **b.** Align those samples to a phylogenetic tree to generate the Matrix as input; **c.** Building and training the ONN prediction model, the ONN prediction model was trained for 30,000 iterations by using samples that have been mapped to the biome ontology; **d.** The ONN prediction model with the highest accuracy on the training set was selected as the final well-trained ONN model; **e.** The well-trained ONN model generates a hierarchical prediction, which indicates the predicted biomes (together with contributions) of the query sample on every layer of the biome ontology.

**Supplementary Figure 3**

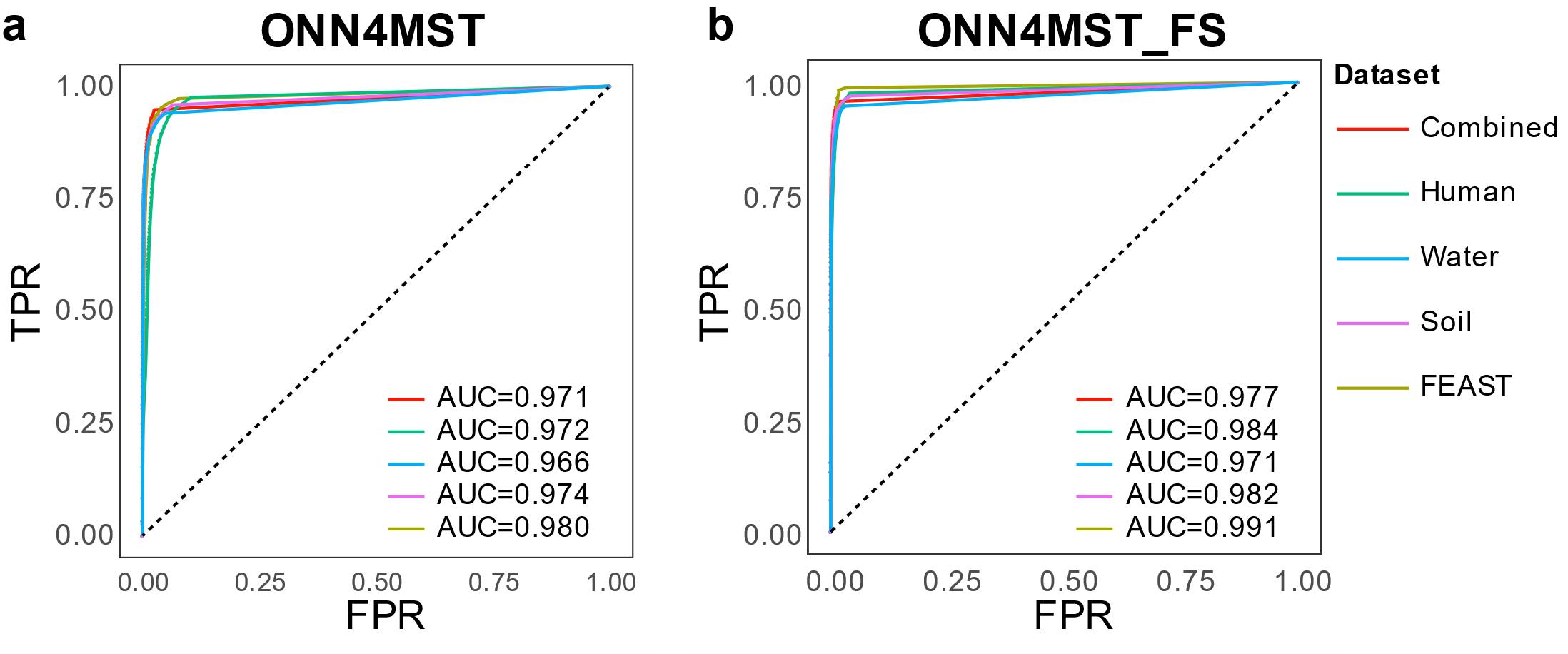

**Supplementary Figure 3. ROC curves using ONN4MST on all five datasets. a.** The ROC curve of ONN4MST using all features on each dataset; **b.** The ROC of ONN4MST using selected features on each dataset; (**Abbreviations**. ONN4MST_FS: ONN4MST using selected features).

**Supplementary Figure 4**

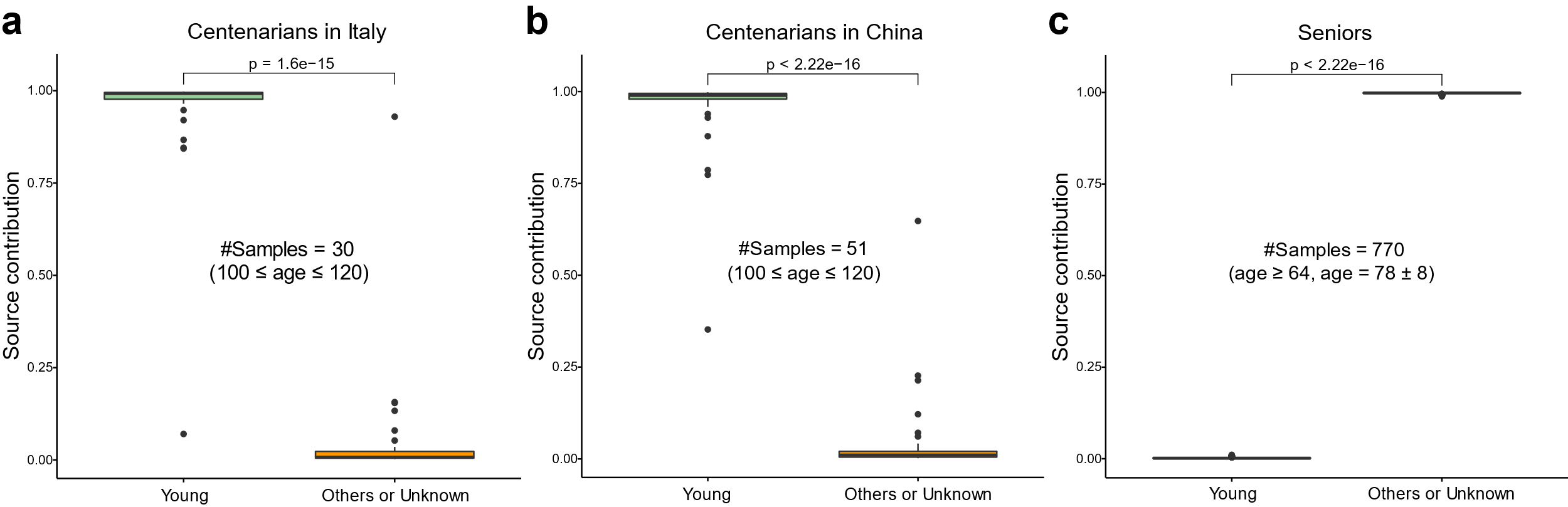

**Supplementary Figure 4. ONN4MST estimations of source contribution to centenarians’ gut microbiome.** **a.** There is a significantly larger “Young human gut” contribution (Wilcoxon-test, p = 1.6e-15) in centenarians from Italy. **b.** There is a significantly larger “Young human gut” contribution (Wilcoxon-test, p <2.22e-16) in centenarians from China. **c.** A large contribution of “Unknown” is assigned in seniors (Wilcoxon-test, p < 2.22e-16).

**Supplementary Figure 5**

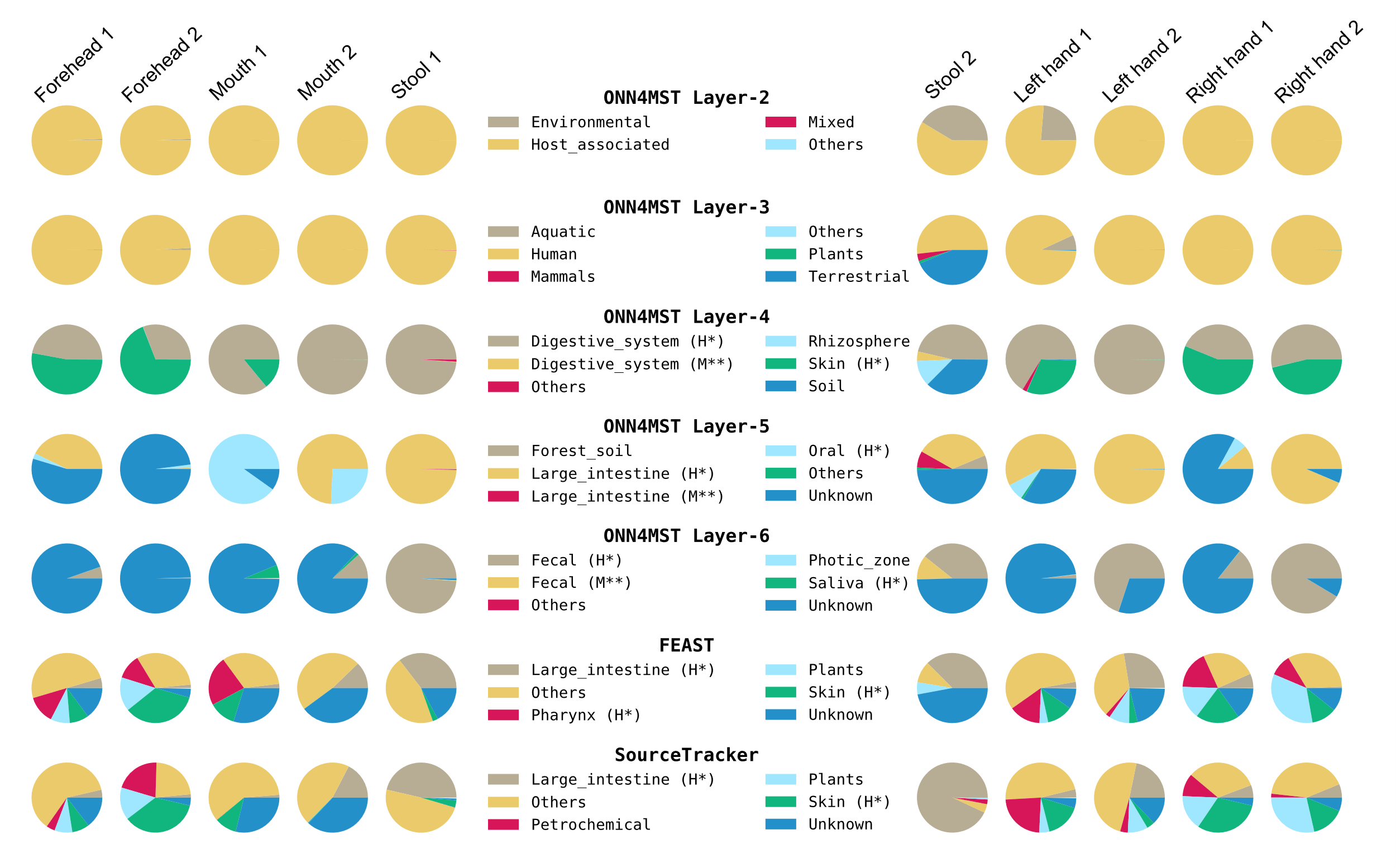

**Supplementary Figure 5. Source tracking of samples from closely related human associated biomes.** ONN4MST could successfully distinguish samples from human forehead, mouth, stool and hands, and could also accurately identify the actual biome for most of these samples. However, FEAST and SourceTracker fall short in identifying these samples as from different human niches, while oral samples were never successfully identified by these two methods.

**Supplementary Figure 6**

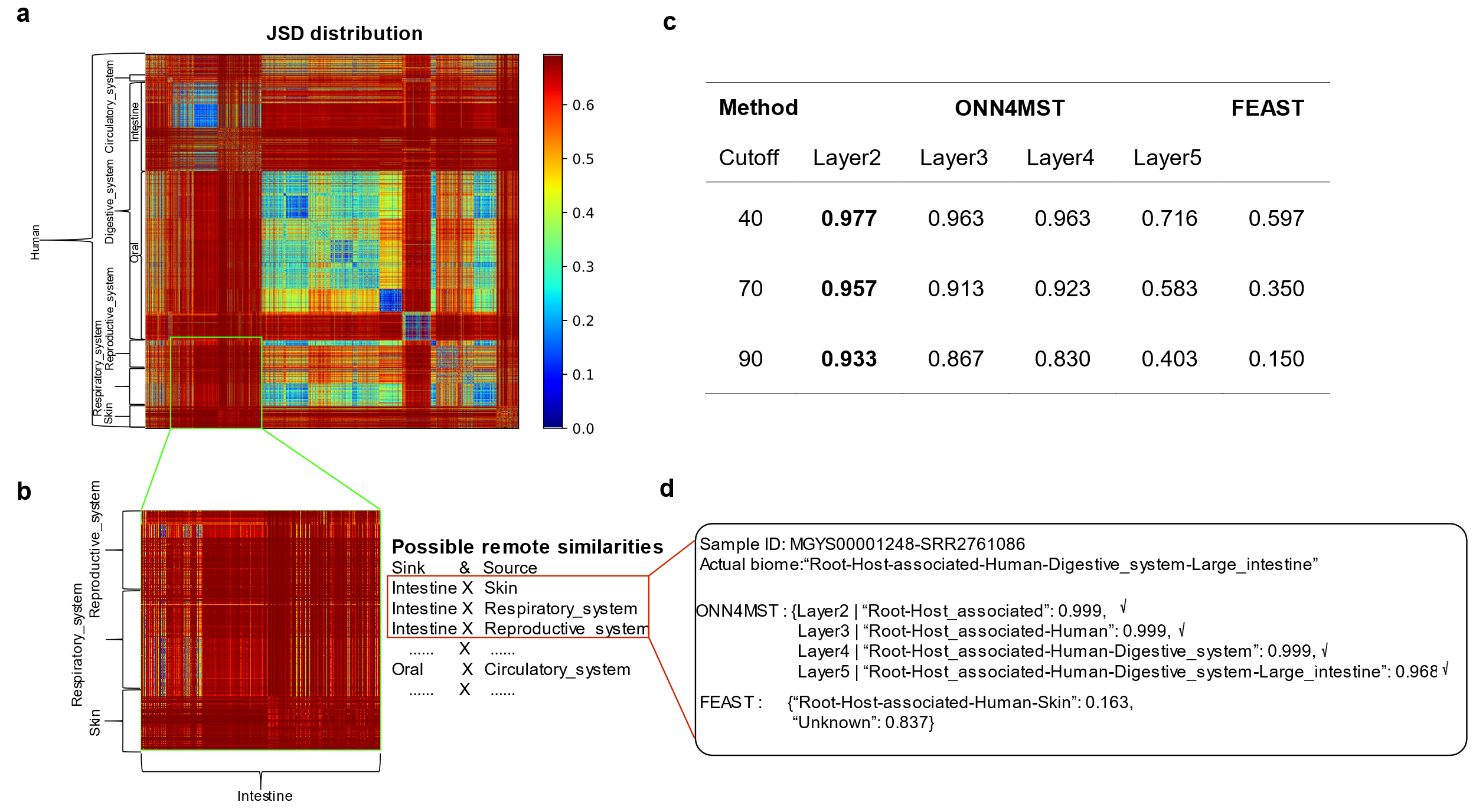

**Supplementary Figure 6: Identification of samples in several ontologically close biomes from Human. a.** Distribution of JSD between samples from “Human” biome; **b.** The distribution of JSD between samples from “Intestine” and “Skin”, “Resporatory_system” and “Reproductive_system”; **c.** The source tracking accuracy by ONN4MST and FEAST with different cutoff (threshold of source contribution); **d.** An example of search result by using ONN4MST and FEAST.

**Supplementary Figure 7**

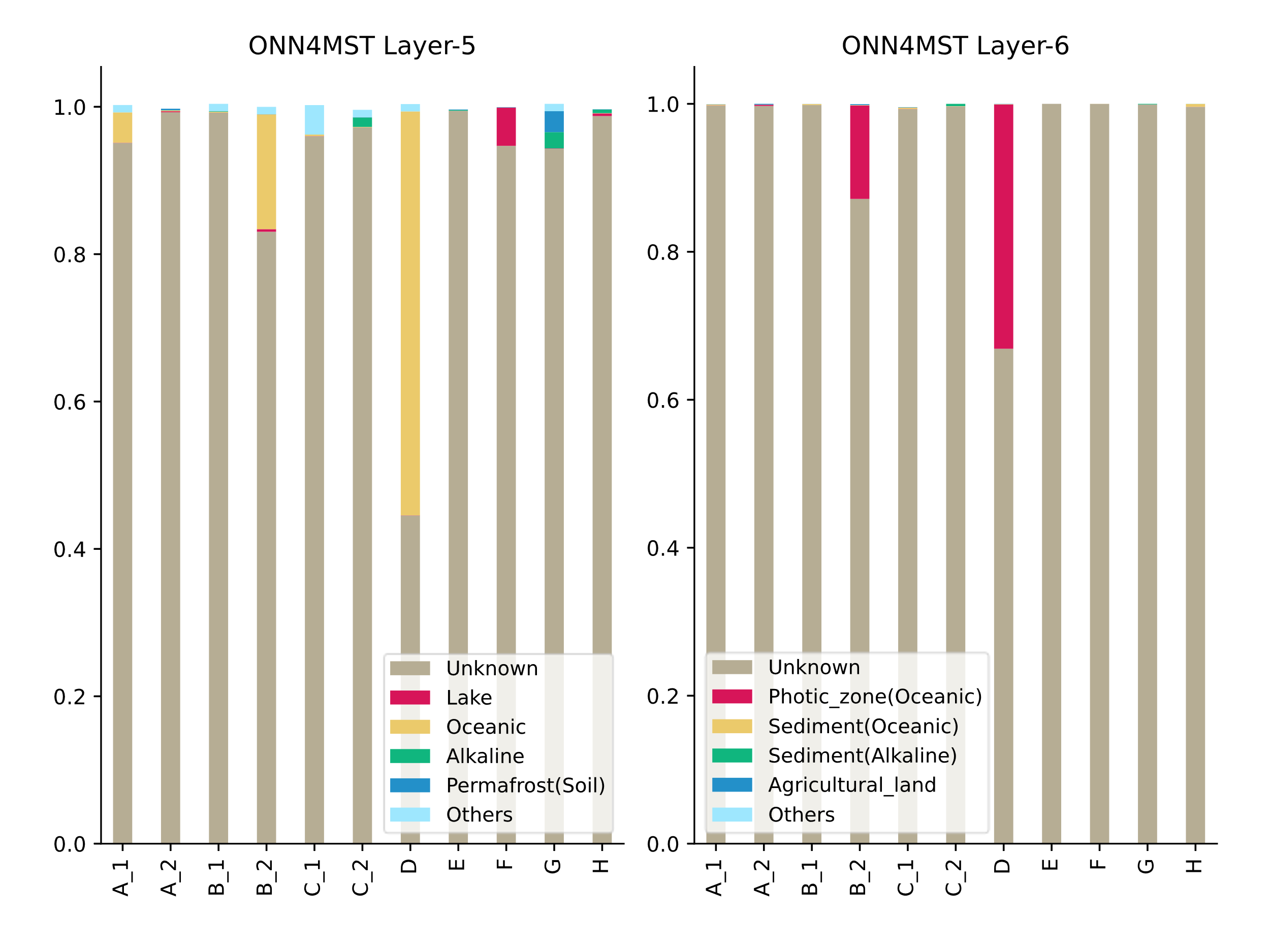

**Supplementary Figure 7. Source tracking results at layer five and layer six for microbiome samples from a less studied biome.** Left: source tracking results at the fifth layer; Right: source tracking results at the sixth layer. Results have shown that ONN4MST could identify the actual source from those polluted microbial communities. A_1, A_2: two samples collected from a single well; B_1, B_2: two samples collected from another single well; C_1, C_2: two samples collected from the third single well; D-H: samples collected from other five wells, respectively.

**Supplementary Figure 8**

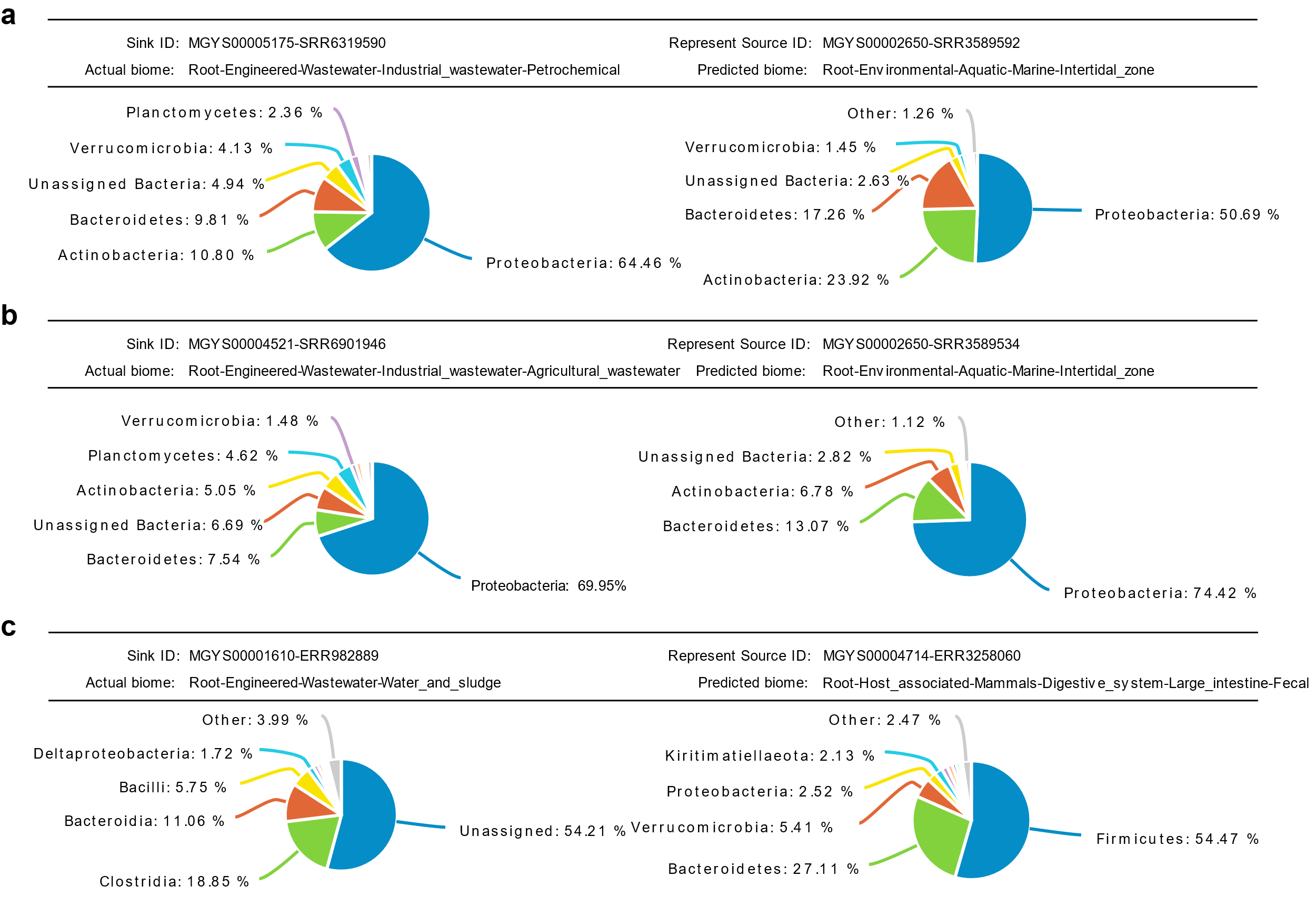

**Supplementary Figure 8. Knowledge discovery of similar samples from ontologically-remote biomes. a.** Sample “MGYS00005175-SRR6319590” (from MGnify database) from biome “Root-Engineered-Wastewater-Industrial_wastewater-Petrochemical” has been identified by ONN4MST as similar with those from biome “Root-Environmental-Aquatic-Marine-Intertidal_zone”; **b.** Sample “MGYS000045215-SRR6901946” (from MGnify database) from biome “Root-Engineered-Wastewater-Industrial_wastewater-Agricultural_wastewater” has been identified by ONN4MST as similar with those from biome “Root-Environmental-Aquatic-Marine-Intertidal_zone”; **c.** Sample “MGYS00001610-ERR982889” (from MGnify database) with annotated biome “Root-Engineered-Wastewater-Water_and_sludge” have been identified by ONN4MST as similar with those from biome “Root-Host_associated-Mammals-Digestive_system-Large_intestine-Fecal”.

**Supplementary Table 1**

**Supplementary Table 1. Microbial community samples and data used for model building and testing.** **Note:** Human*: refers to those samples selected from MGnify biome “Host_associated-Human”. Aquatic**: refers to those samples selected from MGnify biome “Aquatic”. Soil***: refers to those samples selected from MGnify biome “Environmental-Terrestrial” and biome “Plants-Rhizosphere”. FEAST: refers to those samples selected from Shenhav *et al*.^1^. Details are shown in **Supplementary Table 2**, all the data can be downloaded through **Supplementary Table 12**.

| **Dataset** | **Combined** | **Human** | **Water** | **Soil** | **FEAST** |
| --- | --- | --- | --- | --- | --- |
| **Top-level biome** | Root | Human* | Aquatic** | Soil*** | Human gut |
| **Number of biomes involved** | 114 | 25 | 44 | 16 | 3 |
| **Number of samples** | 125,823 | 53,553 | 27,667 | 11,528 | 10,270 |
| **Number of phyla** | 225 | 204 | 222 | 201 | 133 |
| **Number of classes** | 660 | 557 | 653 | 539 | 277 |
| **Number of orders** | 2,018 | 1,412 | 19,85 | 1,394 | 552 |
| **Number of families** | 6,232 | 2,801 | 6,040 | 2,962 | 1,118 |
| **Number of genera** | 16,081 | 6,523 | 15,261 | 6,753 | 3,389 |
| **Number of species** | 45,477 | 16,135 | 36,406 | 12,769 | 5,762 |
| **Average number of species per sample** | 411.22 | 408.55 | 350.23 | 532.12 | 111.05 |
| **Notes** | Selected samples from MGnify | Selected samples from MGnify biome “Human” | Selected samples from MGnify biome “Aquatic” | Selected samples from MGnify biome “Environmental-Terrestrial” and biome “Plants-Rhizosphere” | Selected samples from FEAST dataset^1^ |

**Supplementary Table 2**

**Supplementary Table 2. Biomes and number of samples used in EBI MGnify and this study.** All samples shown in this table belong to the Combined dataset, while certain sets of samples belong to the Human, Water and Soil datasets, as shown in the second column of the table.

| **MGnify biome** | **Belong to** | **ONN4MST biome** | **Number of samples** |
| --- | --- | --- | --- |
| Root |  | Root | 0 |
| Abyssal_plane | Water | Root-Environmental-Aquatic-Marine-Oceanic-Abyssal_plane | 792 |
| Activated_Sludge |  | Root-Engineered-Wastewater-Activated_Sludge | 233 |
| Agricultural | Soil | Root-Environmental-Terrestrial-Soil-Agricultural | 1,132 |
|  | Soil | Root-Environmental-Terrestrial-Soil-Loam-Agricultural | 48 |
| Agricultural_land | Soil | Root-Environmental-Terrestrial-Soil-Crop-Agricultural_land | 16 |
| Agricultural_wastewater |  | Root-Engineered-Wastewater-Industrial_wastewater-Agricultural_wastewater | 34 |
| Alkaline | Water | Root-Environmental-Aquatic-Non_marine_Saline_and_Alkaline-Alkaline | 0 |
| Aphotic_zone | Water | Root-Environmental-Aquatic-Marine-Oceanic-Aphotic_zone | 62 |
| Aquatic | Water | Root-Environmental-Aquatic | 0 |
| Benthic | Water | Root-Environmental-Aquatic-Marine-Oceanic-Benthic | 292 |
| Biofilm | Water | Root-Environmental-Aquatic-Freshwater-Groundwater-Biofilm | 11 |
| Black_smokers | Water | Root-Environmental-Aquatic-Marine-Hydrothermal_vents-Black_smokers | 2 |
| Blood | Human | Root-Host_associated-Human-Circulatory_system-Blood | 24 |
| Boreal_forest | Soil | Root-Environmental-Terrestrial-Soil-Boreal_forest | 3 |
| Buccal_mucosa |  | Root-Host_associated-Mammals-Digestive_system-Oral_cavity-Buccal_mucosa | 26 |
| Cecum |  | Root-Host_associated-Mammals-Digestive_system-Large_intestine-Cecum | 201 |
| Circulatory_system | Human | Root-Host_associated-Human-Circulatory_system | 2 |
| Coastal | Water | Root-Environmental-Aquatic-Marine-Coastal | 4,307 |
| Cold_seeps | Water | Root-Environmental-Aquatic-Marine-Cold_seeps | 555 |
| Contaminated | Soil | Root-Environmental-Terrestrial-Soil-Contaminated | 1 |
| Coral_reef | Water | Root-Environmental-Aquatic-Marine-Intertidal_zone-Coral_reef | 1,004 |
| Crop | Soil | Root-Environmental-Terrestrial-Soil-Crop | 1 |
| Desert | Soil | Root-Environmental-Terrestrial-Soil-Desert | 28 |
| Diffuse_flow | Water | Root-Environmental-Aquatic-Marine-Hydrothermal_vents-Diffuse_flow | 1 |
| Digestive_system |  | Root-Host_associated-Mammals-Digestive_system | 4,941 |
|  | Human | Root-Host_associated-Human-Digestive_system | 1,810 |
|  |  | Root-Host_associated-Insecta-Digestive_system | 165 |
| Dissolved_organics_(aerobic) |  | Root-Engineered-Wastewater-Nutrient_removal-Dissolved_organics_(aerobic) | 48 |
| Dissolved_organics_(anaerobic) |  | Root-Engineered-Wastewater-Nutrient_removal-Dissolved_organics_(anaerobic) | 8 |
| Endophytes |  | Root-Host_associated-Plants-Rhizoplane-Endophytes | 35 |
|  |  | Root-Host_associated-Plants-Phylloplane-Endophytes | 3,025 |
| Engineered |  | Root-Engineered | 0 |
| Environmental |  | Root-Environmental | 0 |
| Epiphytes |  | Root-Host_associated-Plants-Rhizosphere-Epiphytes | 36 |
| Estuary | Water | Root-Environmental-Aquatic-Marine-Intertidal_zone-Estuary | 83 |
| Fecal |  | Root-Host_associated-Mammals-Digestive_system-Fecal | 4,822 |
|  |  | Root-Host_associated-Mammals-Digestive_system-Large_intestine-Fecal | 2,904 |
|  | Human | Root-Host_associated-Human-Digestive_system-Large_intestine-Fecal | 29,427 |
| Female | Human | Root-Host_associated-Human-Reproductive_system-Female | 20 |
| Foregut |  | Root-Host_associated-Mammals-Digestive_system-Foregut | 0 |
| Forest_soil | Soil | Root-Environmental-Terrestrial-Soil-Forest_soil | 230 |
|  | Soil | Root-Host_associated-Plants-Rhizosphere-Forest_soil | 959 |
| Freshwater | Water | Root-Environmental-Aquatic-Freshwater | 9 |
| Grasslands | Soil | Root-Environmental-Terrestrial-Soil-Grasslands | 350 |
| Groundwater | Water | Root-Environmental-Aquatic-Freshwater-Groundwater | 0 |
| Host-associated |  | Root-Host_associated | 0 |
| Human | Human | Root-Host_associated-Human | 1,016 |
| Hydrothermal_vents | Water | Root-Environmental-Aquatic-Marine-Hydrothermal_vents | 52 |
| Hypersaline | Water | Root-Environmental-Aquatic-Non_marine_Saline_and_Alkaline-Hypersaline | 7 |
| Industrial_wastewater |  | Root-Engineered-Wastewater-Industrial_wastewater | 261 |
| Insecta |  | Root-Host_associated-Insecta | 608 |
| Intertidal_zone | Water | Root-Environmental-Aquatic-Marine-Intertidal_zone | 90 |
| Intestine | Human | Root-Host_associated-Human-Digestive_system-Intestine | 553 |
| Lake | Water | Root-Environmental-Aquatic-Freshwater-Lake | 3,475 |
| Large_intestine | Human | Root-Host_associated-Human-Digestive_system-Large_intestine | 535 |
|  |  | Root-Host_associated-Mammals-Digestive_system-Large_intestine | 494 |
| Loam | Soil | Root-Environmental-Terrestrial-Soil-Loam | 0 |
| Lympathic_system | Human | Root-Host_associated-Human-Lympathic_system | 0 |
| Lymph_nodes | Human | Root-Host_associated-Human-Lympathic_system-Lymph_nodes | 12 |
| Mammals |  | Root-Host_associated-Mammals | 0 |
| Mangrove_swamp | Water | Root-Environmental-Aquatic-Marine-Intertidal_zone-Mangrove_swamp | 106 |
| Marginal_Sea | Water | Root-Environmental-Aquatic-Marine-Marginal_Sea | 220 |
| Marine | Water | Root-Environmental-Aquatic-Marine | 5,922 |
| Microbial_mats | Water | Root-Environmental-Aquatic-Marine-Hydrothermal_vents-Microbial_mats | 11 |
| Microbialites | Water | Root-Environmental-Aquatic-Marine-Intertidal_zone-Microbialites | 7 |
| Mine_water |  | Root-Engineered-Wastewater-Industrial_wastewater-Mine_water | 79 |
| Mixed |  | Root-Mixed | 7,231 |
| Nasal_cavity | Human | Root-Host_associated-Human-Respiratory_system-Nasopharyngeal-Nasal_cavity | 17 |
| Nasopharyngeal | Human | Root-Host_associated-Human-Respiratory_system-Nasopharyngeal | 2,637 |
| Neritic_zone | Water | Root-Environmental-Aquatic-Marine-Neritic_zone | 77 |
| Non-marine_Saline_and_Alkaline | Water | Root-Environmental-Aquatic-Non_marine_Saline_and_Alkaline | 0 |
| Nutrient_removal |  | Root-Engineered-Wastewater-Nutrient_removal | 0 |
| Oceanic | Water | Root-Environmental-Aquatic-Marine-Oceanic | 3,457 |
| Oil-contaminated | Water | Root-Environmental-Aquatic-Marine-Oceanic-Oil_contaminated | 69 |
| Oil-contaminated_sediment | Water | Root-Environmental-Aquatic-Marine-Oil_contaminated_sediment | 256 |
| Oil-contaminated_sediments | Water | Root-Environmental-Aquatic-Marine-Oceanic-Oil_contaminated_sediments | 48 |
| Oil_seeps | Water | Root-Environmental-Aquatic-Marine-Oil_seeps | 2 |
| Oral | Human | Root-Host_associated-Human-Digestive_system-Oral | 5,820 |
| Oral_cavity |  | Root-Host_associated-Mammals-Digestive_system-Oral_cavity | 0 |
| Pelagic | Water | Root-Environmental-Aquatic-Marine-Pelagic | 176 |
| Periodontal_pockets | Human | Root-Host_associated-Human-Digestive_system-Oral-Periodontal_pockets | 10 |
| Permafrost | Soil | Root-Environmental-Terrestrial-Soil-Permafrost | 74 |
| Petrochemical |  | Root-Engineered-Wastewater-Industrial_wastewater-Petrochemical | 14 |
| Pharynx | Human | Root-Host_associated-Human-Respiratory_system-Nasopharyngeal-Pharynx | 55 |
| Photic_zone | Water | Root-Environmental-Aquatic-Marine-Oceanic-Photic_zone | 376 |
| Phylloplane |  | Root-Host_associated-Plants-Phylloplane | 21 |
| Plants |  | Root-Host_associated-Plants | 1,986 |
| Pulmonary_system | Human | Root-Host_associated-Human-Respiratory_system-Pulmonary_system | 0 |
| Reproductive_system | Human | Root-Host_associated-Human-Reproductive_system | 0 |
| Respiratory_system | Human | Root-Host_associated-Human-Respiratory_system | 23 |
| Rhizoplane |  | Root-Host_associated-Plants-Rhizoplane | 0 |
| Rhizosphere |  | Root-Host_associated-Plants-Rhizosphere | 3,925 |
| Root |  | Root-Host_associated-Plants-Root | 703 |
| Rumen |  | Root-Host_associated-Mammals-Digestive_system-Foregut-Rumen | 42 |
|  |  | Root-Host_associated-Mammals-Digestive_system-Stomach-Rumen | 612 |
| Saliva | Human | Root-Host_associated-Human-Digestive_system-Oral-Saliva | 3,862 |
| Salt_crystallizer_pond | Water | Root-Environmental-Aquatic-Non_marine_Saline_and_Alkaline-Salt_crystallizer_pond | 108 |
| Salt_marsh | Water | Root-Environmental-Aquatic-Marine-Intertidal_zone-Salt_marsh | 757 |
| Sand | Soil | Root-Environmental-Terrestrial-Soil-Sand | 16 |
| Sediment | Water | Root-Environmental-Aquatic-Marine-Coastal-Sediment | 979 |
|  | Water | Root-Environmental-Aquatic-Sediment | 315 |
|  | Water | Root-Environmental-Aquatic-Marine-Sediment | 3,031 |
|  | Water | Root-Environmental-Aquatic-Marine-Cold_seeps-Sediment | 82 |
|  | Water | Root-Environmental-Aquatic-Non_marine_Saline_and_Alkaline-Alkaline-Sediment | 98 |
|  | Water | Root-Environmental-Aquatic-Marine-Wetlands-Sediment | 105 |
|  | Water | Root-Environmental-Aquatic-Marine-Oceanic-Sediment | 468 |
|  | Water | Root-Environmental-Aquatic-Thermal_springs-Sediment | 10 |
|  | Water | Root-Environmental-Aquatic-Non_marine_Saline_and_Alkaline-Hypersaline-Sediment | 19 |
|  | Water | Root-Environmental-Aquatic-Marine-Neritic_zone-Sediment | 5 |
|  | Water | Root-Environmental-Aquatic-Marine-Intertidal_zone-Sediment | 140 |
|  | Water | Root-Environmental-Aquatic-Marine-Hydrothermal_vents-Sediment | 74 |
| Skin | Human | Root-Host_associated-Human-Skin | 4,848 |
| Soil | Soil | Root-Environmental-Terrestrial-Soil | 5,038 |
|  | Soil | Root-Host_associated-Plants-Rhizosphere-Soil | 3,380 |
| Sputum | Human | Root-Host_associated-Human-Respiratory_system-Pulmonary_system-Sputum | 135 |
| Stomach |  | Root-Host_associated-Mammals-Digestive_system-Stomach | 3 |
| Subgingival_plaque | Human | Root-Host_associated-Human-Digestive_system-Oral-Subgingival_plaque | 794 |
| Supragingival_plaque | Human | Root-Host_associated-Human-Digestive_system-Oral-Supragingival_plaque | 513 |
| Terrestrial |  | Root-Environmental-Terrestrial | 0 |
| Thermal_springs | Water | Root-Environmental-Aquatic-Thermal_springs | 0 |
| Throat | Human | Root-Host_associated-Human-Digestive_system-Oral-Throat | 21 |
| Tropical_rainforest | Soil | Root-Environmental-Terrestrial-Soil-Tropical_rainforest | 191 |
| Uranium_contaminated | Soil | Root-Environmental-Terrestrial-Soil-Uranium_contaminated | 1 |
| Vagina | Human | Root-Host_associated-Human-Reproductive_system-Vagina | 164 |
| Volcanic | Water | Root-Environmental-Aquatic-Marine-Volcanic | 1 |
| Wastewater |  | Root-Engineered-Wastewater | 110 |
| Water_and_sludge |  | Root-Engineered-Wastewater-Water_and_sludge | 508 |
| Wetlands | Soil | Root-Environmental-Terrestrial-Soil-Wetlands | 60 |
|  | Water | Root-Environmental-Aquatic-Marine-Wetlands | 6 |
| buccal_mucosa | Human | Root-Host_associated-Human-Digestive_system-Oral-buccal_mucosa | 289 |
| posterior_fornix | Human | Root-Host_associated-Human-Reproductive_system-Vagina-posterior_fornix | 11 |
| tongue_dorsum | Human | Root-Host_associated-Human-Digestive_system-Oral-tongue_dorsum | 955 |
| **Number of samples in different datasets** | | | |
| Combined | | | 125,823 |
| Human | | | 53,553 |
| Water | | | 27,677 |
| Soil | | | 11,528 |

**Supplementary Table 3**

**Supplementary Table 3. The features used in ONN4MST and the selected features used in ONN4MST_FS.** There are 44,668 taxa (or features) in total used in ONN4MST, while ONN4MST_FS (ONN4MST based on selected features) has utilized only 1,462 selected features (attached in Excel table).

**Supplementary Table 4**

**Supplementary Table 4. Evaluation of ONN4MST using the general model built based on the Combined dataset.**

| **Datasets** | **All features** | | | | | **Selected features** | | | | |
| --- | --- | --- | --- | --- | --- | --- | --- | --- | --- | --- |
|  | Precision | Recall | Accuracy | $F_{max}$ | AUC | Precision | Recall | Accuracy | $F_{max}$ | AUC |
| Human | 0.917 | 0.651 | 0.988 | 0.804 | 0.978 | 0.947 | 0.853 | 0.994 | 0.914 | 0.993 |
| Soil | 0.970 | 0.692 | 0.985 | 0.849 | 0.986 | 0.959 | 0.802 | 0.988 | 0.876 | 0.986 |
| Water | 0.926 | 0.747 | 0.993 | 0.849 | 0.984 | 0.887 | 0.781 | 0.993 | 0.833 | 0.990 |
| FEAST | 0.051 | 0.978 | 0.293 | 0.188 | 0.684 | 0.052 | 0.995 | 0.286 | 0.128 | 0.723 |

**Supplementary Table 5**

**Supplementary Table 5. Evaluation of ONN4MST using the human model trained on the Human dataset.**

| **Datasets** | **All features** | | | | | **Selected features** | | | | |
| --- | --- | --- | --- | --- | --- | --- | --- | --- | --- | --- |
|  | Precision | Recall | Accuracy | $F_{max}$ | AUC | Precision | Recall | Accuracy | $F_{max}$ | AUC |
| Soil | 1 | 0 | 0.895 | 0.19 | 0.5 | 1 | 0 | 0.895 | 0.19 | 0.5 |
| Water | 1 | 0 | 0.976 | 0.047 | 0.5 | 1 | 0 | 0.976 | 0.047 | 0.5 |

**Supplementary Table 6**

**Supplementary Table 6. The results of five biome from “Human” using all features by ONN4MST at layer five.** The results have shown that ONN4MST could still reach a high AUC when the number of samples decrease from five thousand to five hundred.

| **Biome** | **Number of samples** | **Precision** | **Recall** | **Accuracy** | **AUC** |
| --- | --- | --- | --- | --- | --- |
| “Root-Host_associated-Human-Digestive_system-Oral” | 5,820 | 0.932 | 0.965 | 0.990 | 0.996 |
| “Root-Host_associated-Human-Respiratory_system-Nasopharyngeal” | 2,637 | 0.951 | 0.897 | 0.997 | 0.992 |
| “Root-Host_associated-Human-Digestive_system-Large_intestine” | 535 | 0.936 | 0.955 | 0.974 | 0.994 |
| “Root-Host_associated-Human-Reproductive_system-Vagina” | 164 | 0.714 | 0.754 | 0.999 | 0.962 |

**Supplementary Table 7**

**Supplementary Table 7. Running time of different methods when search one query against different datasets. Note:** For N query samples searched against M source samples, since Striped UniFrac only generate a matrix of pair-wise comparison results in a single run, we have added a post-process that converted the running time and memory usage of Striped UniFrac, from the actual time used for generating the matrix of results (M+N)^2^, to the time used for M*N only. This might under-estimate the real time cost by Striped UniFrac for the query.

| **Datasets to be searched** | **ONN4MST** | **ONN4MST_FS** | **FEAST** | **SourceTracker** | **JSD** | **Meta-Prism** | **Striped UniFrac** |
| --- | --- | --- | --- | --- | --- | --- | --- |
| FEAST | 0.13 | 0.034 | 21,876 | 821,066 | 12.93 | 0.03 | 1.62 |
| Soil | 0.14 | 0.037 | 32,077 | 736,789 | 14.72 | 0.04 | 1.81 |
| Water | 0.17 | 0.034 | 113,330 | 1,548,727 | 20.88 | 0.19 | 4.36 |
| Human | 0.19 | 0.033 | 274,066 | 3,692,956 | 49.25 | 0.39 | 8.46 |
| Combined | 0.18 | 0.036 | 481,442 | 7,960,753 | 137.05 | 0.93 | 19.72 |
| 1M | 0.18 | 0.036 | 3,826,343 | 63,269,457 | 1,095 | 7.97 | 200.02 |

**Supplementary Table 8**

**Supplementary Table 8. Running time of different methods when search queries of different sizes against Combined dataset.** **Note:** For N query samples searched against M source samples, since Striped UniFrac only generate a matrix of pair-wise comparison results in a single run, we have added a post-process that converted the running time and memory usage of Striped UniFrac, from the actual time used for generating the matrix of results (M+N)^2^, to the time used for M*N only. This might under-estimate the real time cost by Striped UniFrac for the query.

| **Number of queries** | **ONN4MST** | **ONN4MST_FS** | **FEAST** | **SourceTracker** | **JSD** | **Meta-Prism** | **Striped UniFrac** |
| --- | --- | --- | --- | --- | --- | --- | --- |
| 1 | 0.18 | 0.04 | 481,442 | 7,960,753 | 137.05 | 0.94 | 19.70 |
| 100 | 18.85 | 2.28 | 48,146,202 | 796,080,733 | 14,277 | 93.90 | 1,969.90 |
| 10,000 | 1,726.41 | 120.01 | 4,814,630,598 | 79,608,074,239 | 1,425,029 | 9,388 | 196,985 |
| 1,000,000 | 224,993.90 | 10,177.20 | 481,463,060,786 | 7,960,807,427,834 | 142,500,000 | 938,830 | 19,698,000 |

**Supplementary Table 9**

**Supplementary Table 9.** **Memory utilization of all methods when search one query against different datasets.** **Note:** For N query samples searched against M source samples, since Striped UniFrac only generate a matrix of pair-wise comparison results in a single run, we have added a post-process that converted the running time and memory usage of Striped UniFrac, from the actual time used for generating the matrix of results (M+N)^2^, to the time used for M*N only. This might under-estimate the real time cost by Striped UniFrac for the query.

| **Datasets to be searched** | **ONN4MST** | **ONN4MST_FS** | **FEAST** | **SourceTracker** | **JSD** | **Meta-Prism** | **Striped UniFrac** |
| --- | --- | --- | --- | --- | --- | --- | --- |
| FEAST | 7.58 | 1.72 | 3.02 | 1.14 | 1.97 | 0.13 | 0.01 |
| Soil | 8.14 | 1.73 | 7.20 | 2.17 | 5.91 | 0.14 | 0.01 |
| Water | 7.85 | 1.74 | 18.00 | 4.61 | 11.83 | 0.28 | 0.02 |
| Human | 7.58 | 1.77 | 36.00 | 9.78 | 23.65 | 0.52 | 0.05 |
| Combined | 22.24 | 1.85 | 84.00 | 18.28 | 47.30 | 1.15 | 0.11 |
| 1M | 22.24 | 1.85 | 671.61 | 141.12 | 364.35 | 9.06 | 0.86 |

**Supplementary Table 10**

**Supplementary Table 10. Memory utilization of all methods when search queries of different sizes against Combined dataset.** **Note:** For N query samples searched against M source samples, since Striped UniFrac only generate a matrix of pair-wise comparison results in a single run, we have added a post-process that converted the running time and memory usage of Striped UniFrac, from the actual time used for generating the matrix of results (M+N)^2^, to the time used for M*N only. This might under-estimate the real time cost by Striped UniFrac for the query.

| **Number of queries** | **ONN4MST** | **ONN4MST_FS** | **FEAST** | **SourceTracker** | **JSD** | **Meta-Prism** | **Striped UniFrac** |
| --- | --- | --- | --- | --- | --- | --- | --- |
| 1 | 22.24 | 1.85 | 84.00 | 18.28 | 47.30 | 1.17 | 0.11 |
| 100 | 22.50 | 1.85 | 84.00 | 18.28 | 47.30 | 1.27 | 0.20 |
| 10,000 | 27.32 | 2.07 | 84.00 | 18.28 | 47.30 | 1.46 | 9.48 |
| 1,000,000 | 504.27 | 23.44 | 84.00 | 18.28 | 47.30 | 1.60 | 938.58 |

**Supplementary Table 11**

**Supplementary Table 11. The open searching results by using ONN4MST against the Combined dataset.** Several remotely similar samples among biomes “Engineered”, “Host_associated” and “Environmental” were discovered.

| **Sample ID in Mgnify** | **Actual biome** | **Predicted biome by using ONN4MST** | **Contribution** |
| --- | --- | --- | --- |
| MGYS00005175-SRR6314145 | Root-Engineered-Wastewater-Industrial_wastewater-Petrochemical | Root-Environmental-Aquatic-Marine-Coastal-Sediment | 0.9045 |
| MGYS00005175-SRR6319590 | Root-Engineered-Wastewater-Industrial_wastewater-Petrochemical | Root-Environmental-Aquatic-Marine-Intertidal_zone | 0.9916 |
| MGYS00001653-ERR1201972 | Root-Engineered-Wastewater-Water_and_sludge | Root-Host_associated-Mammals-Digestive_system-Large_intestine-Fecal | 0.9914 |
| MGYS00001652-ERR1201420 | Root-Engineered-Wastewater-Water_and_sludge | Root-Host_associated-Human-Digestive_system-Large_intestine-Fecal | 0.9976 |
| MGYS00001610-ERR982892 | Root-Engineered-Wastewater-Water_and_sludge | Root-Host_associated-Mammals-Digestive_system-Large_intestine-Fecal | 0.9550 |
| MGYS00001652-ERR1201415 | Root-Engineered-Wastewater-Water_and_sludge | Root-Host_associated-Human-Digestive_system-Large_intestine-Fecal | 0.9316 |
| MGYS00001653-ERR1201940 | Root-Engineered-Wastewater-Water_and_sludge | Root-Host_associated-Mammals-Digestive_system-Large_intestine-Fecal | 0.9684 |
| MGYS00001652-ERR1201412 | Root-Engineered-Wastewater-Water_and_sludge | Root-Host_associated-Human-Digestive_system-Large_intestine-Fecal | 0.9957 |
| MGYS00001610-ERR982938 | Root-Engineered-Wastewater-Water_and_sludge | Root-Host_associated-Mammals-Digestive_system-Large_intestine-Fecal | 0.9914 |
| MGYS00001653-ERR1201941 | Root-Engineered-Wastewater-Water_and_sludge | Root-Host_associated-Mammals-Digestive_system-Large_intestine-Fecal | 0.9550 |
| MGYS00001653-ERR1201974 | Root-Engineered-Wastewater-Water_and_sludge | Root-Host_associated-Mammals-Digestive_system-Large_intestine-Fecal | 0.9143 |
| MGYS00001653-ERR1201957 | Root-Engineered-Wastewater-Water_and_sludge | Root-Host_associated-Mammals-Digestive_system-Large_intestine-Fecal | 0.9233 |
| MGYS00001610-ERR982888 | Root-Engineered-Wastewater-Water_and_sludge | Root-Host_associated-Mammals-Digestive_system-Large_intestine-Fecal | 0.9880 |
| MGYS00001610-ERR982933 | Root-Engineered-Wastewater-Water_and_sludge | Root-Host_associated-Mammals-Digestive_system-Large_intestine-Fecal | 0.9720 |
| MGYS00001610-ERR982934 | Root-Engineered-Wastewater-Water_and_sludge | Root-Host_associated-Mammals-Digestive_system-Large_intestine-Fecal | 0.9683 |
| MGYS00001652-ERR1201417 | Root-Engineered-Wastewater-Water_and_sludge | Root-Host_associated-Human-Digestive_system-Large_intestine-Fecal | 0.9982 |
| MGYS00001653-ERR1201939 | Root-Engineered-Wastewater-Water_and_sludge | Root-Host_associated-Mammals-Digestive_system-Large_intestine-Fecal | 0.9792 |
| MGYS00001652-ERR1201410 | Root-Engineered-Wastewater-Water_and_sludge | Root-Host_associated-Human-Digestive_system-Large_intestine-Fecal | 0.9849 |
| MGYS00001610-ERR982890 | Root-Engineered-Wastewater-Water_and_sludge | Root-Host_associated-Mammals-Digestive_system-Large_intestine-Fecal | 0.9792 |
| MGYS00001610-ERR982917 | Root-Engineered-Wastewater-Water_and_sludge | Root-Host_associated-Mammals-Digestive_system-Large_intestine-Fecal | 0.9233 |
| MGYS00001610-ERR982932 | Root-Engineered-Wastewater-Water_and_sludge | Root-Host_associated-Mammals-Digestive_system-Large_intestine-Fecal | 0.9721 |
| MGYS00001610-ERR982940 | Root-Engineered-Wastewater-Water_and_sludge | Root-Host_associated-Mammals-Digestive_system-Large_intestine-Fecal | 0.9143 |
| MGYS00001610-ERR982887 | Root-Engineered-Wastewater-Water_and_sludge | Root-Host_associated-Mammals-Digestive_system-Large_intestine-Fecal | 0.9944 |
| MGYS00001610-ERR982889 | Root-Engineered-Wastewater-Water_and_sludge | Root-Host_associated-Mammals-Digestive_system-Large_intestine-Fecal | 0.9976 |
| MGYS00001610-ERR982898 | Root-Engineered-Wastewater-Water_and_sludge | Root-Host_associated-Mammals-Digestive_system-Large_intestine-Fecal | 0.9130 |
| MGYS00001610-ERR982891 | Root-Engineered-Wastewater-Water_and_sludge | Root-Host_associated-Mammals-Digestive_system-Large_intestine-Fecal | 0.9684 |
| MGYS00001610-ERR982896 | Root-Engineered-Wastewater-Water_and_sludge | Root-Host_associated-Mammals-Digestive_system-Large_intestine-Fecal | 0.9474 |
| MGYS00001586-ERR701617 | Root-Engineered-Wastewater-Industrial_wastewater | Root-Host_associated-Human-Digestive_system-Large_intestine-Fecal | 0.9034 |
| MGYS00001555-ERR380862 | Root-Engineered-Wastewater-Industrial_wastewater | Root-Host_associated-Mammals-Digestive_system-Stomach-Rumen | 0.9334 |

**Supplementary Table 12**

**Supplementary Table 12. Data download links for all five datasets used in this study (attached in Excel table).**

**Supplementary Table 13**

**Supplementary Table 13. Databases and software parameters used in this study.**

| **Database** |  |  |
| --- | --- | --- |
| **Database** | **Author** | **Website** |
| NCBI taxdump (released Feb 1, 2019) | Federhen S, 2012 | https://ftp.ncbi.nlm.nih.gov/pub/taxonomy/taxdump_archive/ |
| MGnify | Mitchell, A. L. *et al*., 2019 | https://www.ebi.ac.uk/metagenomics/ |
| **Software** | | |
| **Software** | **Author** | **Website** |
| textmineR (version 3.0.4) | Tommy Jones | https://github.com/TommyJones/textmineR |
| Striped-Unifrac (committed Aug 1, 2018) | McDonald *et al*., 2018 | https://github.com/biocore/unifrac |
| Meta-Prism (version 2.0) | Mo Zhu, Kai Kang and Kang Ning, 2020 | https://github.com/HUST-NingKang-Lab/Meta-Prism-2.0 |
| SourceTracker (version 1.0) | Knights, D. Kuczynski, J. *et al*., 2011 | https://github.com/danknights/sourcetracker |
| FEAST (committed Aug 30, 2019) | Shenhav, L., Thompson, M. *et al*., 2019 | https://github.com/cozygene/FEAST |
| Python (version 3.7.4) | Oliphant and Travis E, 2007 | https://www.python.org |
| Pandas (version 1.0.1) | McKinney, W. *et al*., 2010 | https://pandas.pydata.org |
| Treelib (version 1.5.5) | Xiaming Chen | https://github.com/caesar0301/treelib |
| Numpy (version 1.16.1) | Stéfan van der Walt *et al*., 2011 | https://numpy.org |
| Scikit-learn (version 0.23.2) | Pedregosa *et al*., 2011 | https://scikit-learn.org |
| Tensorflow (version 1.14.0) | Martín Abadi *et al*., 2015 | https://www.tensorflow.org |
| R (version 3.6.1) | Team, R. C., 2013 | https://www.r-project.org |
| **Parameters** |  |  |
| **Software** | **Parameter** | **Value** |
| RandomForestRegressor | n_estimators | 100 |
|  | random_state | 1 |
|  | max_depth | 10 |
| ONN4MST | activation function | ReLU |
|  | optimizer | Adam |
|  | loss function | Sigmoid_cross_entropy_with_logits |
|  | training batch size | 512 |
|  | learning rate | 1.00E-04 |
|  | iterations | 30,000 |
| FEAST | different_source_flag | 1 |
|  | em_iterations | 1,000 |
| SourceTracker | alpha1 | 0.001 |
|  | alpha2 | 0.001 |
| JSD | coverage | 1,000 |
| Meta-Prism | mode | matrix |
|  | cores | 5 |
| Striped UniFrac | m | weighted_normalized |
|  | n | 30 |

**Supplementary Note 1: Data representation**

Before generating fixed the Matrix for each microbial community sample, we constructed a phylogenetic tree using all taxa involved in the Combined dataset. The phylogenetic tree was used to calculate the relative abundance of taxa in each microbial community sample. A detailed procedure was shown below.

1. Construct phylogenetic tree using all taxa involved in the Combined dataset. Level 2 to level 9 of the phylogenetic tree represent eight taxonomical levels (“sk”, “k”, “p”, “c”, “o”, “f”, “g”, and species “s”), respectively.
2. Map abundance data of taxa to the phylogenetic tree.
3. Update abundance for each taxon in the phylogenetic tree, the abundances of descendants were also added. E.g. the abundance of “sk__Bacteria;k__;p__Actinob
   acteria” was also added to “sk__Bacteria”.
4. Calculate relative abundance for each taxon in the phylogenetic tree.
5. Fill the relative abundance of taxa at seven taxon levels (“sk”, “k”, “p”, “c”, “o”, “f”, and “g”) into the Matrix.
6. Finally, the format of the Matrix is $\left[ \begin{aligned} R_{t_{sk}^{g_{1}}}, R_{t_{k}^{g_{1}}},R_{t_{p}^{g_{1}}},R_{t_{c}^{g_{1}}},R_{t_{o}^{g_{1}}},R_{t_{f}^{g_{1}}},R_{t_{g}^{g_{1}}} \\ R_{t_{sk}^{g_{2}}}, R_{t_{k}^{g_{2}}},R_{t_{p}^{g_{2}}},R_{t_{c}^{g_{2}}},R_{t_{o}^{g_{2}}},R_{t_{f}^{g_{2}}},R_{t_{g}^{g_{2}}} \\ \vdots\vdots\vdots\vdots\vdots\vdots\vdots\\ R_{t_{sk}^{g_{n}}}, R_{t_{k}^{g_{n}}},R_{t_{p}^{g_{n}}},R_{t_{c}^{g_{n}}},R_{t_{o}^{g_{n}}},R_{t_{f}^{g_{n}}},R_{t_{g}^{g_{n}}} \end{aligned} \right]$.

Where $t_{sk}^{g_{n}}$ is a taxon in lineage of $g_{n}$ in the taxonomical level “sk”, $R_{t}$ is relative abundance of taxon $t$.

For instance, for a sample consists of three OTUs with the number of reads 1,639, 240 and 121 assigned respectively to three taxa “sk_Bacteria”, “sk_Bacteria;k_;p_Bacteroidetes”, and “sk_Bacteria;k_;p_Chlamydiae”, their abundances were firstly updated to 2,000 (1,639+240+121), 240, and 121, then normalized to form the Matrix $\left[ \begin{matrix} 1 & 1 & 0.12 & 0 & 0 & 0 & 0 \\ 1 & 1 & 0.06 & 1 & 1 & 1 & 1 \\ \vdots& \vdots& \vdots& \vdots& \vdots& \vdots& \vdots\end{matrix} \right]$, in which 0.06= 121/2,000, and 0.12= 240/2,000. For a detailed description and an example of the data representation, see **Supplementary** **Table 3**.

**Supplementary Note 2: Other source tracking methods used in this study**

Jensen Shannon Divergence^2^ (JSD) is a distance-based approach for comparison of two probability distributions. Firstly, JSD distance was calculated between each pair of microbial community samples. After JSD distances are computed for all pairs of microbial community samples, the normalized average similarity scores (computed using one minus JSD distance) of each microbial community sample to every biome source are regarded as the source contributions.

Striped UniFrac^3^ (Unique Fraction metric) is a phylogenetic-based distance measurement for microbial community samples. It calculates the distance as the fraction of the branch length of the tree that leads to descendants from either one environment or the other, but not both. Striped UniFrac can also be used to produce a distance matrix for the pairwise phylogenetic distances between the sets of samples.

FEAST^1^ is a microbial source tracking method based on the Expected Maximization (EM) algorithm. It processes microbial community samples as follows: the features (abundance of taxa) are firstly normalized, then the FEAST program with default parameters is conducted. Finally, FEAST program outputs source contributions.

SourceTracker^4^ is a microbial source tracking method based on the Bayesian algorithm. It processes microbial community samples as follows: the features (abundance of taxa) are firstly normalized, then the depth of the rarefaction is set to 1,000 when conducting the program. Finally, SourceTracker program outputs source contributions.

Meta-Prism^5^ is microbial community samples searching system, using phylogenetic-based scoring algorithm to calculate similarity score between two microbial community samples. In this study, we have used the program Meta-Prism 2.0 (https://github.com/HUST-NingKang-Lab/Meta-Prism-2.0) for comparison. Firstly, for each pair of microbial samples, Meta-Prism sums up their common component’s proportion based on the phylogenetic relationship of species recursively from each leaf node to the “Root”. After similarity scores are computed for all pairs of samples, normalized similarity score of each sample to each biome source was regarded as source contributions.

**Supplementary Note 3: Source tracking of samples from closely related human associated biomes**

To demonstrate the performance of ONN4MST on distinguishing samples from ontologically close biomes, we have searched 10 human samples from a published metagenome study^6^ (and also from the Human dataset used in this study) against the Combined dataset. Results have shown that ONN4MST could successfully distinguish samples from human forehead, mouth, stool and hands (**Supplementary Fig. 5**), and could also accurately identify the actual biome for most of these samples. However, FEAST and SourceTracker fall short in identifying these samples as from different human niches, while oral samples were never successfully identified by these two methods. Moreover, some ontologically remote biomes have been incorrectly sourced, such as "Plants" and "Petrochemical", by FEAST and SourceTracker. Again, the ontology-aware ability of ONN4MST might be the reason behind: even a large contribution of "Unknown" is still assigned by ONN4MST at the bottom (sixth) layer according to the biome ontology, while at higher layers ONN4MST could correctly assign samples to the biome that is the actual biome or the biomes in which the actual biome belongs to.

Another example is also provided for source tracking a set of samples from biome “Intestine” and “Skin”, with results also showing the superiority of ONN4MST over other methods (**Supplementary Fig. 6**). Thus, ONN4MST is especially useful for differentiating samples from ontologically close biomes, again proving that its results can ensure accurate source tracking.
